# A distinct subset of oligodendrocyte lineage cells interact with the developing dorsal root entry during its genesis

**DOI:** 10.1101/2022.03.10.483786

**Authors:** Lauren A. Green, Robert Gallant, Jacob P. Brandt, Evan L. Nichols, Cody J. Smith

**Author notes:** These authors contributed equally (Lauren A. Green, Robert Gallant). **CORRESPONDING AUTHOR:** Cody J. Smith, PhD, Department of Biological Sciences, University of Notre Dame, 015 Galvin Life Sciences Building, Notre Dame, IN 46556.

## Abstract

Oligodendrocytes are the myelinating cell of the CNS and are critical for the functionality of the nervous system. In the packed CNS, we know distinct profiles of oligodendrocytes are present. Here, we used intravital imaging in zebrafish to identify a distinct oligodendrocyte lineage cell (OLC) that resides on the dorsal root ganglia sensory neurons in the spinal cord. Our profiling of OLC cellular dynamics revealed a distinct cell cluster that interacts with peripheral sensory neurons at the dorsal root entry zone (DREZ). With pharmacological, physical and genetic manipulations, we show that the entry of dorsal root ganglia pioneer axons across the DREZ is important to produce sensory located oligodendrocyte lineage cells. These oligodendrocyte lineage cells on peripherally derived sensory neurons display distinct processes that are stable and do not express *mbp*. Upon their removal, sensory behavior related to the DRG neurons is abolished. Together, these data support the hypothesis that peripheral neurons at the DREZ can also impact oligodendrocyte development.

## INTRODUCTION

Oligodendrocytes are the myelinating glial cell-type of the central nervous system. We know this myelination is essential for the classically described role of insulation that aids in saltatory conductance of neuronal information throughout the body (Nave and Werner, 2014; Allen and Lyons, 2018). Single-cell RNA sequencing approaches have now clearly demonstrated the heterogeneity among such oligodendrocytes (Marques et al., 2016). For example, scRNA sequencing of mature oligodendrocytes (OLs) in the mouse CNS defined 13 distinct populations of mature OLs (Marques et al., 2016). This data, and others, further identifies heterogeneity within subsets of mature OLs enriched in specific regions of the CNS (Marques et al., 2016; Spitzer et al., 2019; Marisca et al., 2020). In zebrafish, immature oligodendrocyte cells have also been described as heterogeneous in the spinal cord (Marisca et al., 2020). While it is now generally accepted that oligodendrocytes display some heterogeneity, the properties of such heterogeneous populations are less understood.

What we do know is that oligodendrocytes heterogeneity in the spinal cord can be driven by distinct progenitors (Richardson et al., 2006). In the spinal cord of mice most OPCs originate from common progenitors in the pMN but a subset of OPCs also arise from *dbx*^+^ cells that are positioned more dorsally in the spinal cord (Spassky et al., 1998; Fogarty, 2005; Kessaris et al., 2006; Richardson et al., 2006; Tripathi et al., 2011; Crawford et al., 2016). The recent model of progenitor recruitment during oligodendrocyte specification, in zebrafish, may update the theory of dorsal vs ventral derived oligodendrocytes, indicating they both may migrate from the pMN domain but at temporally distinct times (Ravanelli and Appel, 2015). These studies may support origin and temporal derivation as drivers of heterogeneity. It is also clear that environmental events that OPCs experience while navigating to their mature location could impact heterogeneity. We know tiling, activity-dependent mechanisms and a plethora of signaling cascades impact oligodendrocyte differentiation, sheath production and survival (Kirby et al., 2006; Emery, 2010; Hughes et al., 2013; Nave and Werner, 2014; Hines et al., 2015; Mensch et al., 2015; Koudelka et al., 2016; Allen and Lyons, 2018). However, these developmental events have mostly been studied without the heterogeneity of oligodendrocytes as a focus.

Here, we utilize time-lapse imaging of zebrafish to characterize a distinct population of oligodendrocyte lineage cells (OLCs) that reside on sensory neurons in the spinal cord. Using this approach, we visualize a subpopulation of OPCs that migrate towards peripheral sensory neurons as their axons enter the spinal cord. These migrating OPCs contact the dorsal root entry zone (DREZ) and immediately migrate dorsally to associate specifically with CNS-located sensory nerves. These sensory OLCs exhibit and maintain unique molecular and process profiles and display a distinct response to OLCs tiling. In the absence of sensory OLCs, we observed an elimination of a behavioral response to a noxious stimulus. The identification of a population of sensory-associated OLCs with anatomical, developmental and functional heterogeneity from classically described oligodendrocytes adds insight to the heterogeneity of oligodendrocyte populations in the nervous system.

## RESULTS

### Profiling of *sox10*^+^-spinal cord cell dynamics reveals a sensory associated OLC

To investigate OLC heterogeneity, we used time-lapse imaging of OLCs in the spinal cord of intact *Tg(sox10:mrfp)* zebrafish(Kirby et al., 2006). We considered that the migration of distinct OL progenitor cells could reveal heterogeneous OL populations. To explore this, we tracked individual oligodendrocyte progenitor cells (OPC) within 40 μm spinal cord z-projections spanning 200 μm in zebrafish from anterior/posterior as they migrated from their precursor domain to when they halted migration to produce oligodendrocyte sheathes. In this profiling, we measured the directional changes and total distance traveled of single oligodendrocytes (Fig. 1A) (n=6 animals, n=20 OLCs). Even these simple two parameters showed at least two clusters of distinct populations of OLCs, one of which includes cells that migrate less than 100 μm with less than 2 directional changes and a larger cluster that migrated with distances that varied from 50-350 μm and greater than 2 directional changes (Fig. 1A) (n=6 animals, n=20 OLCs).

**Figure 1.**
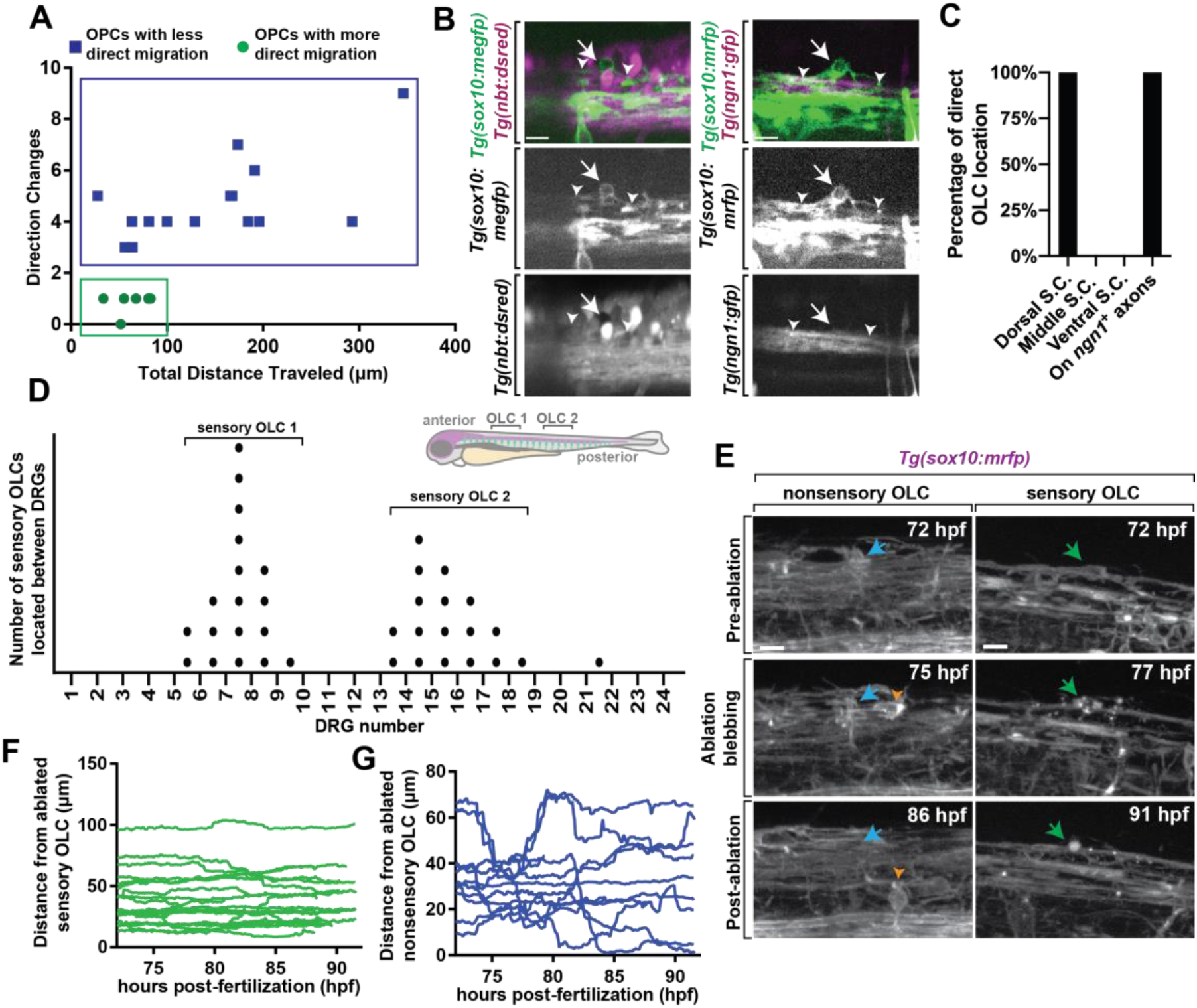
A distinct population of OPCs associates with sensory nerves. (A) Plot of individual OPCs with less direct migration versus individual OPCs with a more direct migration. Green dots denote OPC that migrate less and directly. Blue dots denote OPC with longer distances of migration. (B) Confocal z-stack images from *Tg(sox10:megfp);Tg(nbt:dsred)* and *Tg(ngn1:gfp);Tg(sox10:mrfp)* animals at 72 hpf showing that dorsal migrating oligodendrocytes associate specifically with sensory neurons. White arrow represents oligodendrocyte associated with sensory neurons and white arrowheads represent sensory axons. (C) Quantification of the percentage of OLCs located across specific regions in the spinal cord. (D) Quantification of the number of sensory OLs located between specific DRGs. Each point represents one individual sensory OL. Zebrafish diagram in the upper right represents locations of OLC 1 and 2 alonge the spinal cord. (E) Images from a 24 hour time-lapse starting at 72 hpf in *Tg(sox10:mrfp)* animals showing the ablation of sensory and nonsensory OLs and surrounding OPC response. Blue arrow represents a nonsensory OL. Green arrow represents a sensory OL. Orange arrowheads represent a responding OPC following nonsensory OL ablation. (F) Quantification of the distance surrounding OPCs traveled over time immediately following sensory OL ablation. (G) Quantification of the distance surrounding OPCs traveled over time immediately following nonsensory ablation. Scale bar equals 10μm (B,E).

To determine if these distinct migration patterns dictate potential heterogeneity of OLs we determined if such clusters corresponded with specific anatomically distinct populations. We noted that a cluster of migrating cells resided on the dorsal side of the spinal cord. To determine if they are associated with a specific neuronal population we marked subsets of neurons with specific transgenes. We first scored the location of the OLCs in *Tg(sox10:megfp); Tg(nbt:dsred)* animals which use regulatory sequences of *sox10* to mark oligodendrocytes and their progenitors and *nbt* to panneuronally label axons (Fig. 1B,C)(n=19 animals total)(Peri and Nüsslein-Volhard, 2008; Smith et al., 2014). In this analysis, 100% of the direct-migration cluster resided on dorsally located *nbt*^+^ tracts, indicating their association with neurons (Fig. 1C)(n=10 animals with *nbt*). The location of these dorsal OLCs corresponded near the dorsolateral fasciculus which contains Rohon Beard sensory neurons and DRG sensory neurons in zebrafish. To test if they resided on DRG sensory neurons we similarly scored their association in *Tg(ngn1:gfp); Tg(sox10:mrfp)* animals (Kirby et al., 2006; Prendergast et al., 2012), which mark DRG sensory neurons at 72 hpf with *ngn1*^+^; *sox10*^+^. Rohon Beard neurons, in contrast, do not express *sox10* (Fig. 1B,C). 100% of the direct-migration cluster generated OLCs that reside on the DRG sensory neurons (Fig. 1C) (n=9 animals with *ngn1*). Throughout the manuscript, we define sensory OLCs as *sox10*^+^ cells that reside on the DRG sensory neurons and non-sensory OLCs as *sox10*^+^ cells that are not located on the sensory neurons.

We further explored the anatomical positioning of sensory OLCs by scoring their location in the spinal cord. To do this, we created images that tiled the entirety of the right side of the spinal cord in *Tg(sox10:mrfp);Tg(ngn1:gfp)* animals, allowing us to identify the sensory neurons and the sensory associated OLCs. In this analysis, sensory associated OLCs were distinctly concentrated in two locations in the zebrafish spinal cord. These locations corresponded with DRGs 5-10 and 13-18, with each area having one sensory associated OLC per side of the spinal cord (Fig. 1D) (n=9 animals). All our analysis below was completed at DRGs 6-10 to maximize our likelihood of visualizing these cells. Collectively, these data led us to further characterize at least one small subpopulation of anatomically distinct OLCs.

### Sensory OLCs are not replaced by surrounding OPC populations following ablation

It is well documented that OPCs intrinsically replenish damaged or apoptotic OLs to fill the newly vacant space following OL death (Kirby et al., 2006; Hughes et al., 2013). We therefore next asked if surrounding nonsensory OLC populations could replace an ablated sensory OLC. To test this, we ablated individual sensory OLCs in *Tg(sox10:mrfp)* animals and imaged surrounding OL and OPC response from 72-96 hpf (Fig. 1E)(n=32 animals)(Green et al., 2019). As a control for this, we also ablated individual nonsensory OLCs and imaged the surrounding OL response from 72-96 hpf. In cases of sensory OLC ablation, no surrounding *sox10*^+^ cell responded to the site of ablation (Fig. 1F)(n=19 animals). In cases of nonsensory OL ablation, surrounding *sox10*^+^ cells immediately responded to the site of ablation as previously reported (Fig. 1G)(n=13 animals). These data demonstrate a difference between the sensory subpopulation of OLCs and non-sensory associated OLCs, as they are not replaced by surrounding OLCs.

### Sensory OLCs have distinct sheathes

We next tested if these sensory OLCs have other distinct structural properties by first measuring their cellular processes. We compared these processes to other classically described OLCs. To investigate this, we imaged sensory OLCs in *Tg(nkx2.2a:gfp);Tg(sox10:mrfp)* animals at 72 hpf and measured the sheath width and sheath length of both sensory and classically-described nonsensory OLCs (Fig. 2A)(Kirby et al., 2006). The average process length of sensory OLC was 187.78 ± 15.20 μm compared to an average of 94.67 ± 9.88 μm (p<0.0001) for nonsensory OLs, indicating sensory OLCs exhibit longer and thinner processes/sheathes than nonsensory OLs (Fig. 2B) (n=12 sensory OLC, 11 nonsensory OLC, p>0.0001). Additionally, the average width of sensory OLCs process was 5.01 ± 0.23 μm compared to an average of 7.49 ± 0.41 μm for nonsensory OLs, demonstrating that sensory OLCs have a distinct process profile compared to other nonsensory OLs (Fig. 2C) (n=12 sensory OLC, 11 nonsensory OLC, p>0.0001).

**Figure 2.**
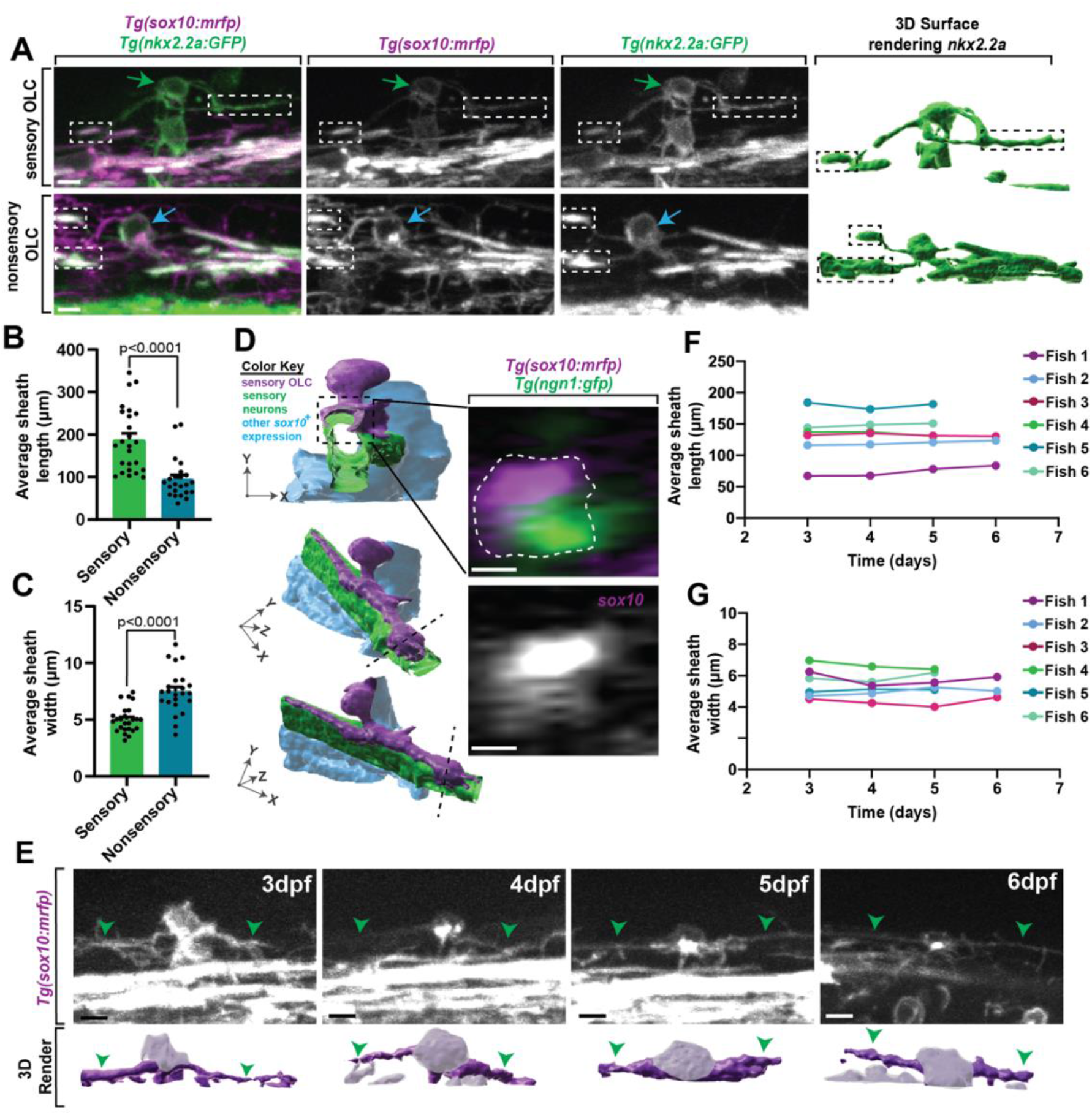
Sensory OLs maintain a distinct sheath profile. (A) Confocal z-stack still images of *Tg(nkx2.2a:gfp);Tg(sox10:mrfp)* animals at 72 hpf showing the distinct sheath profile of sensory OLs compared to nonsensory OLs. Green arrows represent a sensory OL. Blue arrows represent a nonsensory OL. White and black boxes represent examples of sheathes. (B) Quantification of the average sheath length of sensory OLs compared to nonsensory OLs (p<0.0001). (C) Quantification of the average sheath width of sensory OLs compared to nonsensory OLs (p<0.0001). (D) 3D IMARIS surface rendering (left) of a sensory OLC in *Tg(sox10:mrfp);Tg(ngn1:gfp)* animals from a confocal z-stack image (right) represented at three different angles to show ensheathment of sensory axons. White dashed line represents the cross section view. (E). Confocal z-stack images (top row) and IMARIS 3D surface rendering (bottom row) of an individual sensory OL in the same *Tg(sox10:mrfp)* animal at 3dpf, 4dpf, 5dpf, and 6dpf (top row). Green arrowheads represent the sensory axon. Grayed out regions on surface rendering represent sensory OL cell body. (F) Quantification of the average sheath length of the same OL in the same animal over time. (G) Quantification of the average sheath width of the same OL in the same animals over time. Scale bar equals 10μm (A, D-E).

Oligodendrocytes ensheath axonal domains whereas precursor and progenitor populations do not (Nave and Werner, 2014; Allen and Lyons, 2018). To determine if these unique processes ensheath, similar to other oligodendrocyte populations, we imaged *Tg(ngn1:gfp); Tg(sox10:mrfp)* animals at 3 dpf. These animals label sensory neurons in GFP and mRFP and sensory OLC in mRFP. We then reconstructed the surfaces of the cells and digitally rotated the images. In these images, mRFP cell membrane can be visualized outside and ensheathed around GFP; mRFP axons, likely indicating ensheathment of the axon. These data indicate the sensory OLC at 3 dpf have at least initiated ensheathment of axons, likely designating them as oligodendrocytes, albeit by current definitions (Fig. 2D). Furthermore, time-lapse imaging with *Tg(sox10:mrfp)* animals revealed that oligodendrocyte lineage cells located on sensory axons do not divide, at least during our 24 hour imaging windows, potentially arguing again against them representing an immature progenitor or precursor cell.

It is possible that the difference in processes was an indication of immaturity, where sensory-associated OLC represent less mature cells. In zebrafish, classically-described oligodendrocytes do not change the number of myelin sheathes after the first 3 hours of ensheathment(Watkins et al., 2008; Czopka et al., 2013). We therefore asked if this distinct cellular process profile in sensory-associated OLCs is maintained over time. To test this, we located one sensory OLC in individual *Tg(nkx2.2a:gfp);Tg(sox10:mrfp)* animals at 3 dpf. We then imaged that same sensory OLC per animal at 4 dpf, 5 dpf, and 6 dpf, far exceeding the published 3-hour window of sheath dynamics (Fig. 2E)(n=6 animals)(Watkins et al., 2008; Czopka et al., 2013). To ask if individual cellular processes themselves were stable, we measured the length and width of the individual processes. These measurements showed the average process length per sensory OLC per animal was maintained from 3 dpf to 6 dpf (Fig. 2F) (fish 1 p=0.8621, fish 2 p=0.9969, fish 3 p=0.9988, fish 4 p=0.9217, fish 5 p=0.9604, fish 6 p=0.9562). Similarly, these measurements also showed the average process width per sensory OLC per animal was maintained from 3 dpf to 6 dpf (Fig. 2G) (fish 1 p=0.7275, fish 2 p=0.9834, fish 3 p=0.9207, fish 4 p=0.9375, fish 5 p=0.9890, fish 6 p=0.9574). While the length and width of the sheathes did not change overtime, a flattening of the cell body could be visualized (Fig. 2E). Overall, this data shows that sensory OLCs have a cell-process profile that is maintained over time.

### Sensory associated OLCs are produced from the same spinal domain as other OLCs

To uncover potential events that impact sensory OLCs production we first considered the possibility that sensory OLCs were produced from different oligodendrocyte precursors or progenitors that precedes their arrival to the sensory neurons. In the spinal cord, current literature states that the majority of oligodendrocytes are produced from the *olig2*^+^ pMN with a subset potentially produced from a more dorsal *dbx*^+^ domain (Richardson et al., 2006). To determine if these cells are derived from progenitors from different domains we used transgenic markers consistent with different regions of the spinal cord (Fig. S1A). In this analysis, 100% of the sensory associated OLCs expressed *Tg(sox10:mrfp)* and *Tg(sox10:megfp)* that label OPCs and oligodendrocyte (Fig. S1B) (n=25 animals total). *Tg(dbx:gfp)* labeled 0% of sensory-associated OLCs, indicating that these sensory oligodendrocytes unlikely directly originated from the *dbx^+^* region of the spinal cord (n=7 animals)(Briona and Dorsky, 2014). *Tg(nkx2.2a:gfp)* labeled 50% of the sensory-associated OLC indicating a subset originated from the area around the floor plate region of the spinal cord (n=8 animals). All the sensory-associated OLCs were labeled with *Tg(olig2:dsred)* specifying they originated from the pMN domain of the spinal cord like other oligodendrocyte populations (n=7 animals)(Park et al., 2002a). These data, in conjunction with previous literature, support the conclusion that both sensory and non-sensory OLC are derived from the same spinal domain.

If these sensory OLCs are derived from the pMN domain like other oligodendrocytes, then their migration should originate from those ventral spinal domains. We tested this possibility with a second complementary assay by photoconverting OPCs in the ventral spinal region. To trace the direct migration of a single cell to the sensory axon from the ventral spinal cord region, we used *Tg(sox10:eos)* to photoconvert all ventral OPCs and imaged migration from 48-72 hpf (Fig. S1C) (n=3 animals)(McGraw et al., 2012; Green and Smith, 2018). We observed photoconverted (red) cells migrate dorsally from the ventral spinal cord directly to the sensory axon (Fig. S1D). These data are inconsistent with the conclusion that the sensory OLCs are a result of production from different spinal cord precursor/progenitor domains, at least defined by current spinal domain literature.

### Sensory OLCs interact with the dorsal root entry zone

We next considered the hypothesis that their migration exposed them to unique spatiotemporal experience. We noted in timelapse movies that these directed OPCs migrated dorsally and halted their migration in the dorsal spinal cord, close to the dorsal root entry zone (DREZ)(Smith et al., 2017). We therefore asked if this cluster of OPCs contact the DREZ during migration. To test this, we imaged *Tg(sox10:gal4;uas:lifeact-gfp)* zebrafish which allow us to visualize thin projections from the OPCs and sensory neurons during pioneer axon entry(Hines et al., 2015; Nichols and Smith, 2019a, 2019b; Zhang et al., 2019). Since axon entry occurs between 48-72 hpf we also quantified the number of OPCs located on sensory nerves from 36-96 hpf (Fig 3A,B) (n=20 animals). No OPCs were present on sensory axons from 36-48 hpf, however an average of 0.57 ± 0.291 OPCs were present on sensory axons from 48-72 hpf and an average of 1.2 ± 0.202 OPCs were present from 72-96 hpf (Fig. 3A,B) (n=20 animals). Our data further indicate that 80% of OPCs in the directed cluster contacted the DREZ before they halted their migration and produced sheathes on sensory axons (Fig. 3C) (n=6 animals). After DREZ contact, these OPCs migrated an average of 17.10 μm dorsally until they halted further migration (Fig. 3D) (n=6 animals).

**Figure 3.**
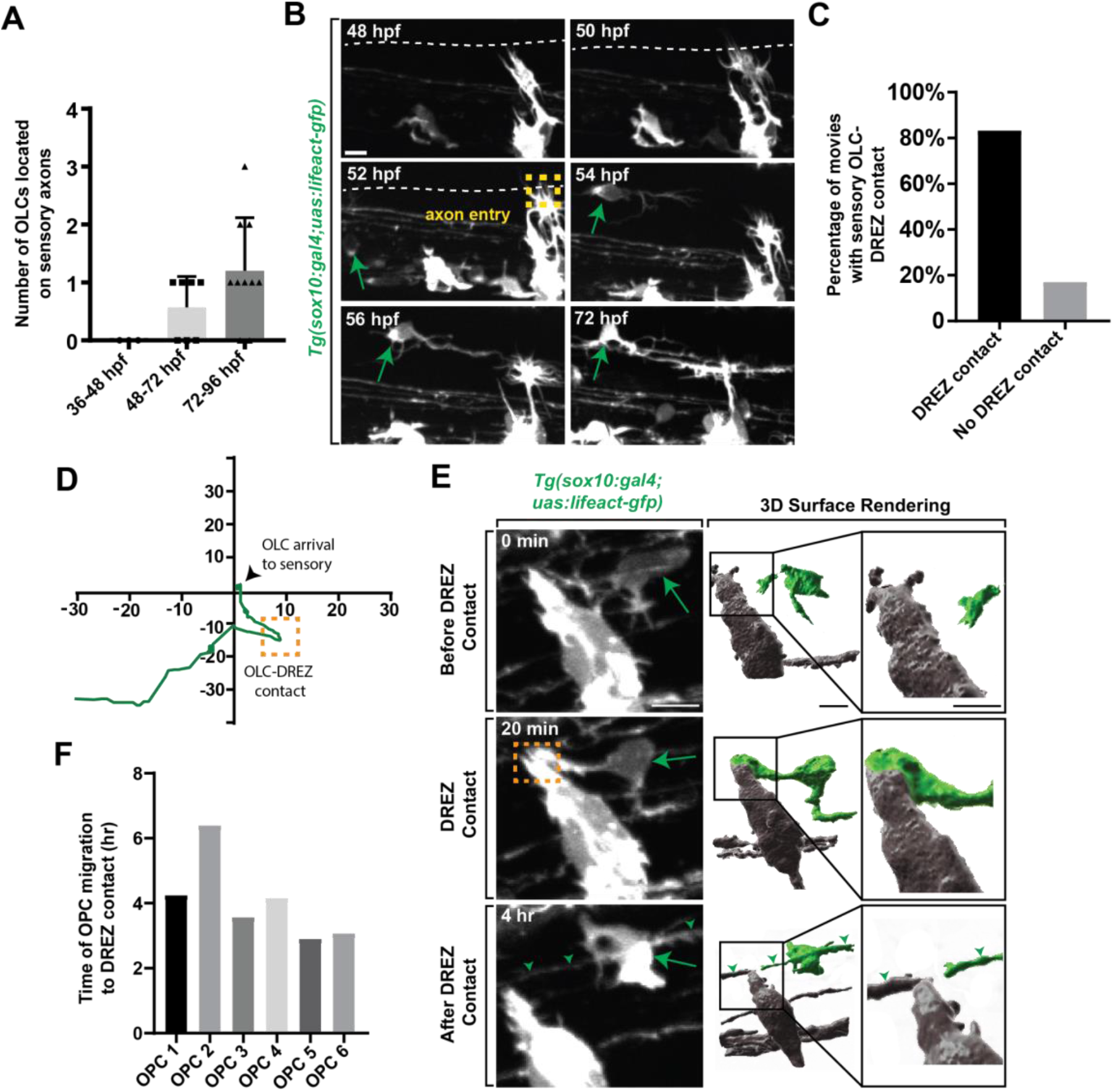
OPCs contact the DREZ immediately following axon entry. (A) Quantification of the number of OPCs located with DRG sensory nerves from 36-96 hpf. (B) Images from a 24 hour time-lapse movie starting at 48 hpf in *Tg(sox10:gal4; uas:lifeact-gfp)* zebrafish showing OPC migration initiation following axon entry. Green arrow represents migrating OPC. Yellow box represents axon entry. (C) Quantification of the percentage of movies where oligodendrocytes arrived to the sensory with or without DREZ contact. (D) Migration plot of the path an oligodendrocyte traveled from the ventral spinal cord region to sensory axons. Orange box represents DREZ contact. (E) Images from a 24 hour time-lapse movie starting at 48 hpf in *Tg(sox10:gal4; uas:lifeact-gfp)* zebrafish showing an oligodendrocyte extending a cellular process that contacts the DREZ. Green arrow represents OPC projection that contacts the DREZ. Orange box indicates OPC-DREZ contact. Green arrowheads represent the sensory axon. IMARIS 3D surface rendering of the 20 min confocal image in showing an OLC cellular process contacting the DREZ. Grey cells are DRG and peripherally located. Green cell is the sensory OL. (F) Quantification of the time it takes for an individual OPC to initiate migration post-axon entry and contact the DREZ. Scale bar equals 10μm (B,E).

Unlike other events that have been identified to promote oligodendrocyte development, the DREZ is composed of peripheral nervous system components(Golding et al., 1997; Smith et al., 2017), making it somewhat surprising that it could influence CNS development. To begin to address the role of the DREZ in sensory OLCs we assessed when the DREZ-directed OPCs contact the DREZ. For this study, we define the genesis of the DREZ as the time in development when the peripheral nervous system derived pioneer DRG axons enter the spinal cord from their peripheral location(Nichols and Smith, 2019a). In zebrafish, this occurs between 48-72 hpf. The precise spatiotemporal location of pioneer axon entry and DREZ genesis can be identified by the formation of actin-based invasive structures in pioneer axons (Nichols and Smith, 2019a, 2019b; Zhang et al., 2019; Kikel-Coury et al., 2021). To ask if DREZ-directed OPCs contact the DREZ during this process we collected z-stacks every 5 mins from 48-72 hpf in *Tg(sox10:gal4; uas:lifeact-GFP)* animals (Fig. 3E,F) (n=6 animals). In these animals, both oligodendrocytes and sensory neurons are labeled with Lifeact-GFP but separated spatially, with OLCs in the spinal cord and sensory neurons in the periphery. In these movies, 100% of the DREZ-directed OPCs contacted the DREZ after the axons had entered the spinal cord, contacting the peripheral portion of the cells at the DREZ (Fig. 3E, Fig. S2A) (n=6 animals). Collectively in the movies, OPC DREZ-contact occurred 4.06 hours after pioneer axon entry at the DREZ (Fig. 3F) (Average time of OPC migration to DREZ per OPC: OPC 1 = 4.25, OPC 2 = 6.40, OPC 3 = 3.58, OPC 4 = 4.16, OPC 5 = 2.91, OPC 6 = 3.08). These results led us to the hypothesis that formation of the DREZ could impact the migration of OPCs from their ventral progenitor.

### Formation of the DREZ is required to produce sensory OLCs

If the genesis of the DREZ directs sensory OLCs, then perturbing DREZ formation should alter the production of these sensory OLCs. To test this, we altered DREZ formation by disrupting DRG pioneer axons from entering the spinal cord as previously reported (Nichols and Smith, 2019a, 2019b; Zhang et al., 2019). We first did this by blocking DRG pioneer axon entry components using SU6656, an inhibitor of Src that is required for pioneer axons to enter the spinal cord(Nichols and Smith, 2019a). In this experiment *Tg(sox10:mrfp); Tg(ngn1:gfp)* animals were treated with SU6656 or DMSO from 36-72 hpf to block axon entry (Fig. 4A,B) (n=27 animals total)(Nichols and Smith, 2019a). We imaged a spinal region in these animals that spanned DRGs at somites 6-10 in which we know sensory OLCs are located and scored the number of sensory OLCs present (Fig 4C) (n=8 animals). In DMSO-treated spinal segments, an average of 0.875 ± 0.085 sensory OLCs were present by 72 hpf (n=16 animals). This was in contrast to an average of 0.091 ± 0.090 sensory OLs that were present in SU6656-treated spinal segments where DREZ formation was perturbed (n=11 animals) (Fig. 4C, p<0.0001). These results support the conclusion that failure to form the DREZ could result in reduced migration of OPCs to the sensory axons.

**Figure 4.**
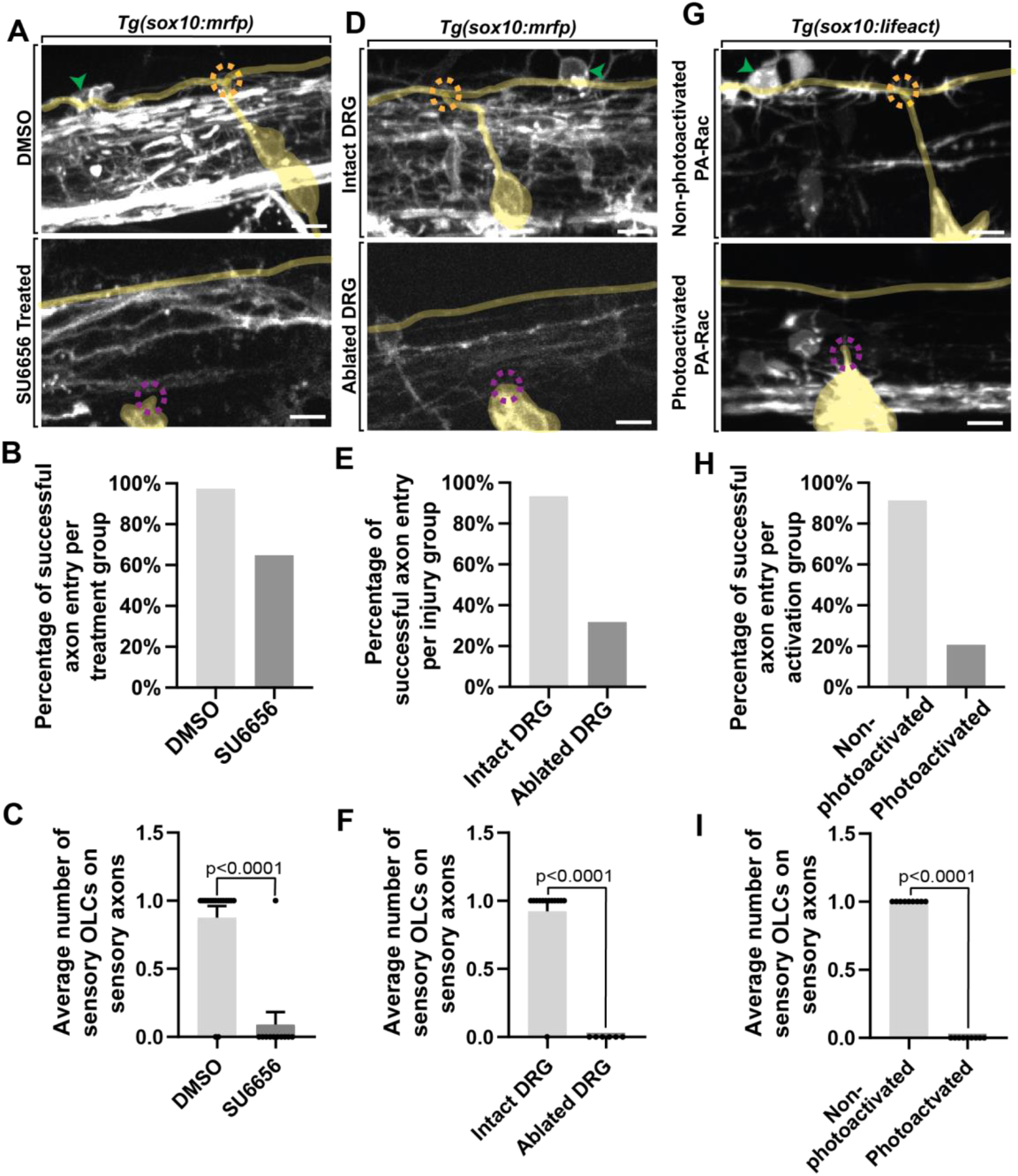
Failed axon entry does not result in sensory OLCs. (A) Confocal z-stack images of *Tg(sox10:mrfp);Tg(ngn1:gfp)* zebrafish at 3 dpf showing axon entry and the presence or absence of a sensory OLC in DMSO control animals compared to SU6656 treated animals. (B) Quantification representing the percentage of successful axon entry events per DMSO group versus SU6656 treatments. (C) Quantification of the average number of sensory OLCs on sensory axons in DMSO versus SU6656 treated animals (p<0.0001). (D) Confocal z-stack images of *Tg(sox10:mrfp); Tg(ngn1:gfp)* zebrafish at 3 dpf showing axon entry and the presence or absence of a sensory OL in intact animals compared to animals with ablated DRGs. (E) Quantification representing the percentage of successful axon entry events per intact group versus ablated DRGs. (F) Quantification of the average number of sensory OCLs on sensory axons in intact DRGs versus ablated DRGs (p<0.0001). (G) Confocal z-stack images of *Tg(sox10:gal4; uas:lifeact-gfp)* animals injected with PA-Rac1 at 3 dpf showing axon entry and the presence or absence of a sensory OLC in non-photoactivated animals compared to photoactivated animals. (H) Quantification representing the percentage of successful axon entry events per non-photoactivated group versus the photoactivated group. (I) Quantification of the average number of sensory OLCs on sensory axons in non-photoactivated versus photoactivated animals (p<0.0001). All green arrowheads represent sensory OLCs. All dashed orange circles represent successful axon entry. All dashed magenta circles represent failed axon entry. Yellow tracing overlays highlight the DRG and sensory axons (A,D,G). Scale bar equals 10μm (A,D,G).

Since this pharmacological treatment is not cell-specific, we also used complementary physical manipulations to test the requirement of DREZ formation to drive sensory OLCs. In this paradigm, pioneer axon entry was prevented by ablating the DRG neurons before they produce their pioneer axons and imaged as in our pharmacological experiments (Fig. 4D,E) (n=19 animals total). In control spinal segments where the DRG pioneer neuron was not ablated, an average of 0.93 ± 0.077 sensory OLCs were present at intact spinal segments (n=13 animals). This was a stark contrast to spinal segments where the pioneer neurons were ablated, in which we could not detect any sensory OLCs (n=6 animals) (Fig. 4F, p<0.0001). We could identify sensory-associated OLCs in DRG ablated animals because adjacent regions to the four-ablated nerves still entered and produced spinal projections.

We extended this analysis by altering the DREZ formation with a third complementary approach. Rac1 activation drives the disassembly of invasive structures and thus by mosaically expressing photoactivatable Rac1 in pioneer neurons and photo activating them during DREZ formation, we can block entry of pioneer axons into the spinal cord (Nichols and Smith, 2019a; Zhang et al., 2019). We injected animals with photoactivatable Rac1 protein at the single-cell stage and imaged *Tg(uas:lifeact-gfp); Tg(sox10:gal4) uas:PA-Rac-mcherry* animals from 48-72 hpf (Fig. 4G,H) (n=18 animals total)(Nichols and Smith, 2019a; Zhang et al., 2019; Kikel-Coury et al., 2021). Mosaic animals that were selected for this experiment contained PA-Rac-mcherry in the DRG neurons but not sensory OLCs, allowing us to manipulate the sensory ingrowth without impacting OLCs directly with Rac1 (Fig. 4I). In control animals where the PA-Rac protein was injected, but not photoactivated with 445 nm, an average of 1.0 ± 0.111 sensory OLCs could be detected in spinal segments (n=9 animals). In contrast, sensory OLCs could not be identified in spinal segments with photoactivated Rac1 (n=9 animals) (Fig. 4I, p<0.0001). The simplest explanation for the data from these three manipulations of DREZ formation is consistent with the possibility that DREZ formation is important to produce sensory associated OLCs.

If the formation of the DREZ drives production of distinct OLCs, then altering the timing of DREZ formation should also change the timing of DREZ-directed OPCs and formation of sensory OLCs. To address this possibility, we induced early entry of pioneer axons by forcing the formation of invasive structures in pioneer DRG axons as previously noted (Fig. 5A) (n=8 animals). To do this, we treated *Tg(sox10:gal4; uas:lifeact-gfp)* animals with paclitaxel, a potent activator of invasive structures, from 36-72 hpf, then imaged them from 48-72 hpf (Fig. 5A, C,D) (n=8 animals)(Nichols and Smith, 2019a, 2019b; Kikel-Coury et al., 2021). To first confirm that pioneer axons entered the spinal cord early we scored the timing of pioneer axon entry. We normalized these time points in DMSO and Paclitaxel treated animals by scoring the time of entry compared to the initiation of the pioneer axon. Consistent with previous reports (Nichols and Smith, 2019a, 2019b; Kikel-Coury et al., 2021), paclitaxel-treated animals had early genesis of the DREZ of 2.33 ± 0.439 hours on average compared to DMSO-treated animals where DREZ genesis occurred at 4.48 ± 0.338 hours on average (Fig. 5C, p=0.0081) (n=8 animals). In support of the hypothesis that DREZ formation drives the production of sensory OLCs, paclitaxel-treated animals formed DREZ-directed OPCs at 0.71 ± 0.217 hours compared to DMSO that formed at 5.71 ± 1.322 hours on average (Fig. 5D,G) (p=0.0338, n=8 animals). As an argument against the possibility that this accelerated OPC migration was a result of increased velocity of OPCs, we tracked individual DREZ-directed OPCs as they migrated. The velocity of OPC migration in DMSO treated animals was 0.15 ± 0.037 on average compared to 0.08 ± 0.030 in paclitaxel-treated animals (Fig. S2B, p=0.3001, n=8 animals). The duration of time between initiation of OPC migration and arrival to the DREZ in DMSO-treated animals was 4.50 ± 0.936 hours on average compared to 1.58 ± 0.096 hours in paclitaxel-treated animals (Fig. S2C, p=0.0710, n=8 animals). Both the velocity and time of OPC migration were statistically indistinguishable. Together, these data again support the possibility that the DREZ is important to produce sensory-associated OLCs.

**Figure 5.**
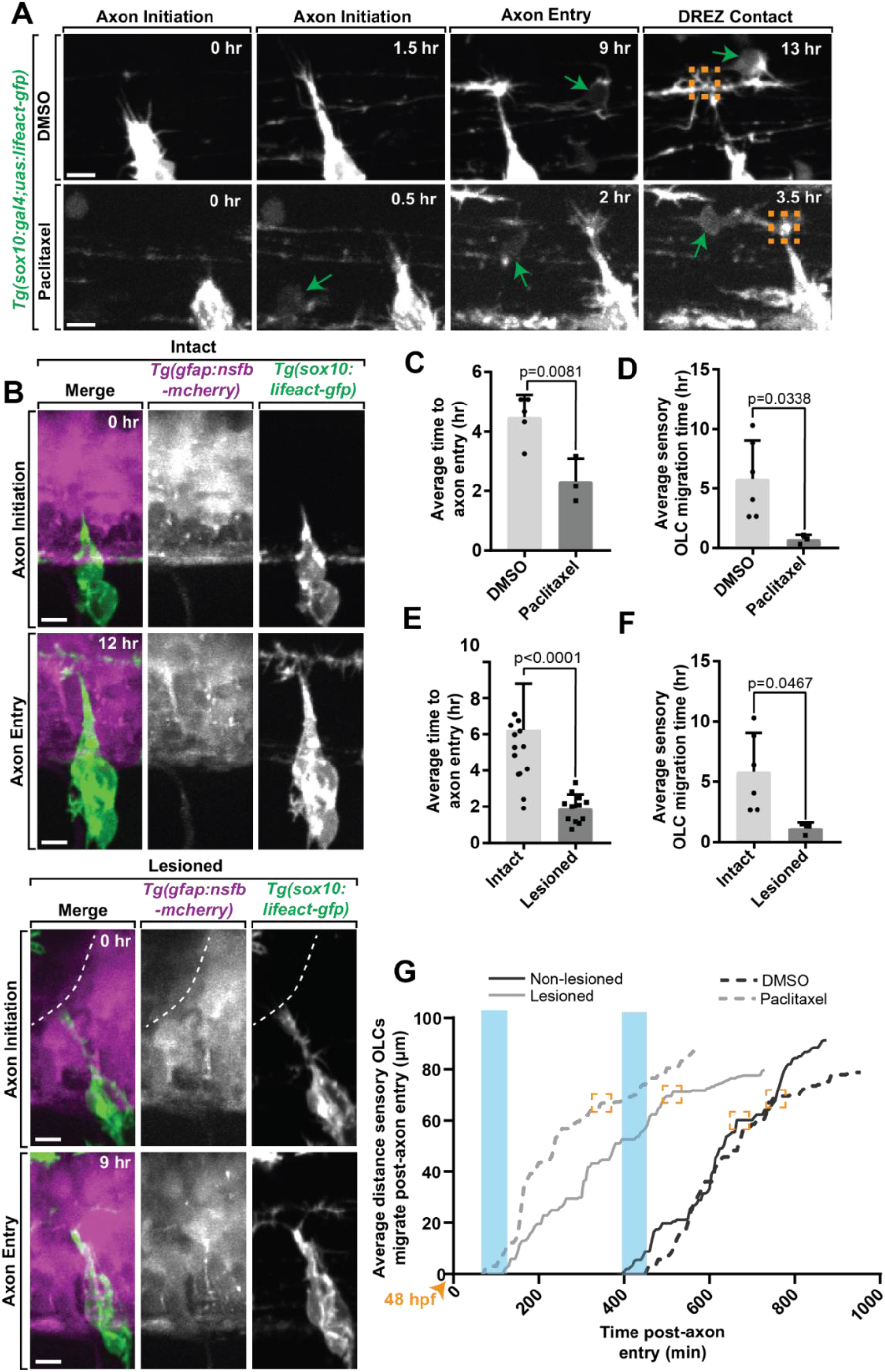
Early axon entry promotes early OPC migration to sensory nerves. (A) Images from a 24 hour time-lapse movie starting at 48 hpf in *Tg(sox10:gal4 : uas:lifeact-gfp)* zebrafish treated with Paclitaxel showing axon initiation and sensory OLC and DREZ contact earlier than typical axon entry. Green arrows represent migrating sensory OLC. (B) Still Images from a 24 hour time-lapse movie starting at 48 hpf in intact and lesioned *Tg(sox10:gal4;uas:lifeact-gfp);Tg(gfap:nsfb-mcherry)* zebrafish also showing axon initiation and successful axon entry earlier than typical axon entry. White dashed line indicates lesion site. (C) Quantification of average time to axon entry in DMSO and Paclitaxel treated animals (p=0.0081). (D) Quantification of the average time to sensory OLC migration post-axon entry in DMSO and Paclitaxel treated animals (p=0.0338). (E-F) Paralleled quantifications of C-D in Lesioned and Non-lesioned animals (D = p<0.0001, E = p=0.0467). (G) Quantification of the average distance and time three OLCs migrated post-axon entry in DMSO and Paclitaxel treated animals and lesioned and non-lesioned animals. Orange arrowhead indicates start of timelapse at 48 hpf. Blue rectangles highlight the time of axon entry. Orange boxes indicate OLC-DREZ contact. Scale bar equals 10μm (A-B).

We next considered the possibility that the peripherally-derived sensory pioneer axon itself could provide a cue to drive sensory OLCs. Alternatively, the actual act of the sensory axon breaching the radial glial limitans to cross into the spinal cord could drive the formation of these sensory OLCs. To provide insight into these two possibilities, we sought to mimic this potential breach by creating a break in the radial glial limitans with a physical manipulation (Nichols and Smith, 2019a). In this paradigm, we created a lesion of the radial glial limitans in a 10-12 μm region of the spinal directly dorsal to the motor exit point where the DREZ consistently forms, prior to pioneer axon initiation in *Tg(sox10:megfp); Tg(gfap:mcherry)* animals at 48 hpf(Nichols and Smith, 2019a). After ablation, we imaged the radial glia lesion site from 48-72 hpf and quantified the time of axon initiation to axon entry, or DREZ formation in lesioned and non-lesioned animals (Fig. 5B,E-G) (n=9 animals total). On average, axons enter in non-lesioned animals at 6.24 ± 0.712 hours compared to 1.90 ± 0.225 hours in lesioned animals (Fig. 5E, p<0.0001). We then compared the timing of sensory OLC formation and DREZ contact of the OPC. Tracing DREZ directed OPC-cells in these movies revealed that creating an artificial breach of the spinal cord induced OPC-directed migration at an average of 2.0 ± 0.282 hr compared to non-lesioned regions that migrated at an average of 6.0 ± 1.322 hrs (Fig. 5F, p=0.0467). Arguing against the possibility that this expedited OPC migration was from increased velocity of the OPCs in the control vs experimental paradigm, the time for OPC migration from its initiation to the DREZ or velocity during that time was not significantly different (Fig. S2D-E, p=0.5592 and p=0.5445). Collectively, we observed accelerated OPC initiation of migration was in both Paclitaxol-treated and physically-breached animals compared to DMSO-treated and non-breached animals (Fig. 5G). The simplest explanation for this data is that the act of pioneer axon entry impacts DREZ-directed OPCs.

### Sensory OLCs have distinct molecular properties to non-sensory OLCs

Given the distinct sheath profile and development of sensory OLCs, we next asked whether these cells express myelin-associated factors distinct of other mature myelinating oligodendrocyte (Nave and Werner, 2014). To test this, we imaged *Tg(mbp:gfp); Tg(sox10:mrfp)* animals at 5 dpf and 15 dpf and scored the number of sensory-associated OLCs that were *mbp^+^* (Fig. 6A) (n=55 animals total)(Koudelka et al., 2016). We observed 100% of sensory OLCs at 5 dpf animals were *sox10*^+^, but 0% were *mbp*^−^ at 5 dpf (Fig. 6B, n=9 5 dpf animals). To argue against the possibility that these cells presented immature *mbp*^+^ cells, we also imaged the same *Tg(mbp:gfp); Tg(sox10:mrfp)* animal at 5 dpf and 15 dpf, which again extends past the typical differentiation period of an oligodendrocyte in zebrafish (Czopka et al., 2013). At 15 dpf, we observed 100% sensory OLCs that were *sox10*^+^ and 0% sensory OLCs that were *mbp*^+^, matching the *mbp* expression in 5 dpf sensory OLCs (Fig. 6B, n=9 15 dpf animals). We also quantified the marked *mbp* expression in nonsensory OLCs (Fig. 6C, n=18 animals total). We observed 25% of nonsensory OLCs were *mbp*^−^ while 75% of nonsensory OLCs were *mbp*^+^ at 5 dpf (n=9 5 dpf animals). Similarly at 15 dpf we observed 30% of nonsensory OLCs were *mbp*^−^ while 70% of nonsensory OLCs were *mbp*^+^ (n=9 15 dpf animals). To test that the lack of *mbp* expression in sensory OLCs was not specific to the transgene, we also stained animals at 15 dpf with anti-MBP (Fig. 6D). At 15 dpf, there were an average of 0.0 ± 0 sensory OLCs expressing MBP^+^ (per imaging window) compared to an average of 6.2 ±1.2 nonsensory OLCs that were MBP^+^ (per imaging window)(Fig. 6E, n=5 animals total, p=0.0009). These data are consistent with the possibility that sensory OLCs do not express myelin-associated factors at 5 dpf or 15 dpf, past the typical initiation period of ensheathment (Watkins et al., 2008; Czopka et al., 2013). It is important to note that although they do not express myelination components, it is unlikely that these cells represent NG2 cells because of their expression of *nkx2.2*, which NG2 cells do not express(Hughes and Appel, 2016).

**Figure 6.**
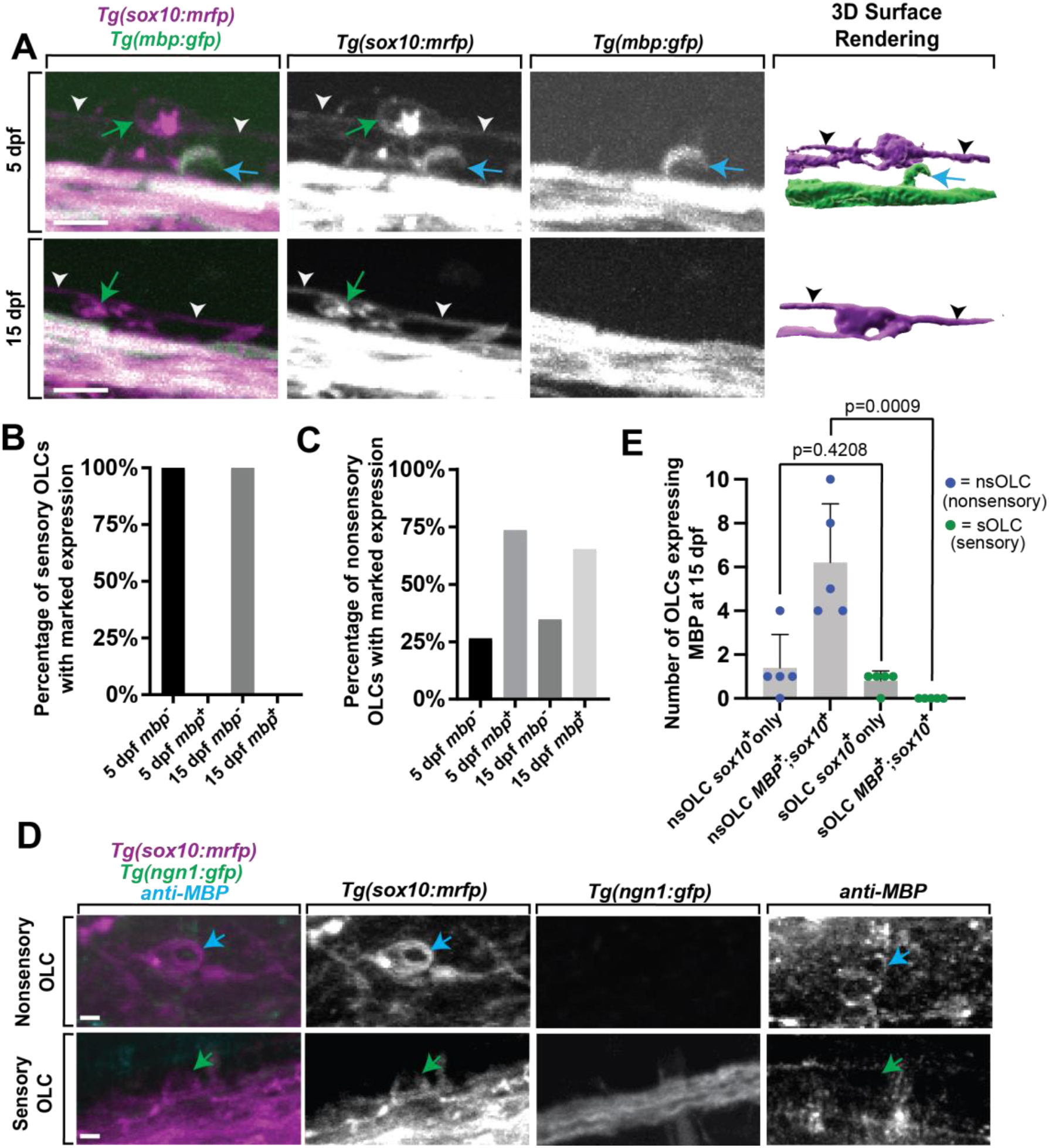
Sensory OLCs do not express typical OLC *mbp* expression. (A) Confocal z-stack still images of *Tg(mbp:gfp);Tg(sox10:mrfp)* animals at 5dpf and 15dpf showing the lack of *mbp*^+^ expression in sensory oligodendrocytes. Blue arrows represent nonsensory OLCs. Green arrows represent sensory OLCs. White and black arrowheads indicate sensory axons. (B) Quantification of the percentage of *mbp*^−^;*sox10*^+^ and *mbp*^+^;*sox10*^+^ sensory OLs at 5dpf and 15dpf. (C) Quantification of the percentage of *mbp*^−^;*sox10*^+^ and *mbp*^+^;*sox10*^+^ nonsensory OLs at 5dpf and 15dpf. (D) Confocal z-stack still images of *Tg(sox10:mrfp);Tg(ngn1:gfp)* animals at 15dpf stained with an anti-MBP antibody showing the lack of anti-MBP in sensory OLCs. Blue arrows represent nonsensory OLCs and green arrows represent sensory OLCs. (E) Quantification of the number of nonsensory OLCs (nsOLCs) and sensory OLCs (sOLCs) expressing *sox10*^+^ only or at 15 dpf. Scale bar equals 10μm (A,E).

### Sensory OLCs display distinct Ca^2+^ transients that non-sensory OLCs

Oligodendrocytes have been shown to have both spontaneous and evoked Ca^2+^ transients. We next investigated such transients in sensory OLCs. DRG neurons can respond to a multitude of stimuli(Lumpkin and Caterina, 2007) and previous literature indicated DRG neurons could be active after exposure to 4°C (Fosque et al., 2015; Nichols and Smith, 2020). We first tested this by imaging *Tg(neurod:gal4); Tg(uas:gCaMP6s)* and measured calcium transients in DRG neurons after exposure to 4°C or 23°C water at 3 dpf (n=21 DRG cells). *Tg(neurod:gal4); Tg(uas:gCaMP6s)* animals were anesthetized for mounting purposes then transferred back into 23°C water without anesthesia before the beginning of the assay (n=5 animals). Unanesthetized animals were imaged at two second intervals, imaged for one minute, exposed to 23°C water, and then imaged for an additional one minute, after which 4°C water was added followed by another one minute of imaging (Fig. 7A) (n=21 DRG cells). In this analysis, the Z score of DRG neuron integral densities were measured and used to assess significant changes in gCaMP6s intensity. If gCaMP6s integral density Z score was 2 or greater, this was marked as an active firing event. (Fig. 7B) (n=5 animals). To verify that this activation was consistent across multiple DRG, we created heat maps of each DRG neuron and their change in gCaMP6s intensity Z score at each timepoint (Fig. 7C) (n=5 animals). Using a strict threshold to create a binary assessment of DRG activation, animals exposed to 4°C water resulted in a significant change in the percent of active DRG neurons at an average of 96% as compared to 6.6% activation in 23°C water (Fig. 7D, p<0.0001, n=5 animals). These results are consistent with the hypothesis that zebrafish DRG neurons respond to 4°C.

**Figure 7.**
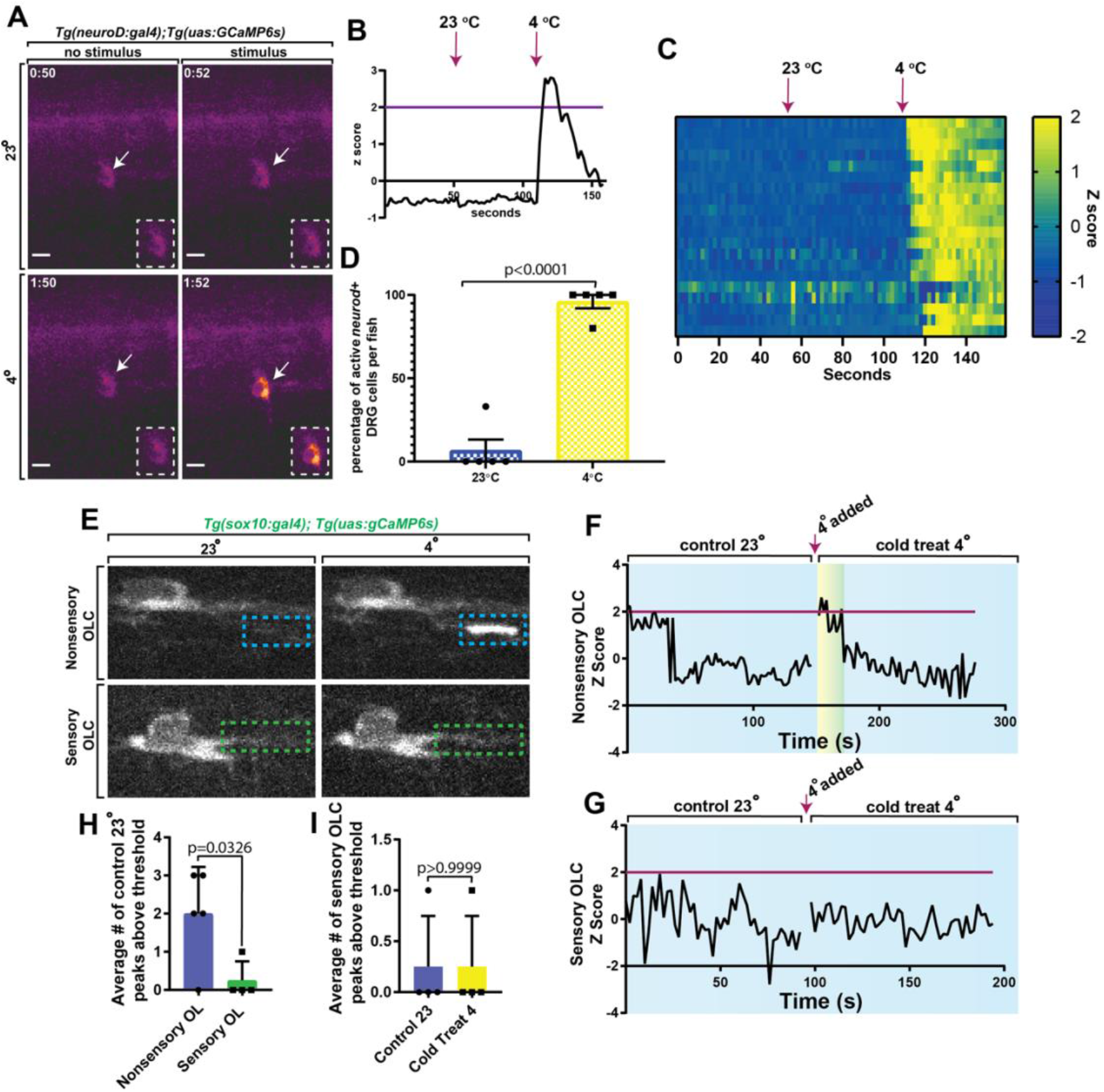
Sensory OLCs display distinct Ca^2+^ transients compared to non-sensory OLCs. (A) Images of *Tg(neurod:gal4); Tg(uas:gCaMP6s)* at 3 dpf showing calcium transients in DRG neurons after exposure to 23°C or 4°C water. (B) Quantification of the Z Score of gCaMP6s intensity over time. Magenta line indicates Z score threshold above 2. Magenta arrow indicates the point at which 23°C or 4°C water was added. (C) Heatmap representing each DRG neuron and the relative change in gCaMP6s intensity between exposure of 23°C or 4°C water over time. (D) Quantification of the percentage of active DRG *neuroD*^+^ cells per animal in 23°C control temperature compared to 4°C water (p<0.0001). (E) Images from a 3 minute time-lapse in *Tg(sox10:gal4); Tg(uas:gCaMP6s)* zebrafish showing calcium transients in nonsensory OLCs compared to sensory OLCs upon exposure to 4°C water. Blue dashed boxes indicate the region of Ca^2+^ transient activity in nonsensory OLCs. Green dashed boxes indicate lack of Ca^2+^ transient activity in sensory OLCs. (F) Quantification of Z Score representing Ca^2+^ transient activity in nonsensory OLCs upon exposure to 4°C water. (G) Quantification of Z Score representing the lack of Ca^2++^ transient activity in sensory OLCs upon exposure to 4°C water. Magenta line indicates standard deviation threshold above 2 and magenta arrow indicates the point at which 4°C water was added in both F and G. Any peak above this magenta line indicates a positive peak value for calcium expression. (H) Quantification of the average number of control 23°C peaks above threshold in sensory OLCs versus nonsensory OLCs (p=0.0326). (I) Quantification of the average number of sensory OLCs peaks above the threshold in control 23°C versus 4°C water animals (p > 0.9999). Scale bar equals 10 μm (A,E)

We therefore next tested if sensory OLCs display Ca^2+^ characteristics like non-sensory OLCs in both spontaneous and evoked scenarios (Krasnow et al., 2018; Marisca et al., 2020). We first asked if sensory OLCs display spontaneous activity like those described in previous reports (Hines et al., 2015; Mensch et al., 2015; Baraban et al., 2017; Krasnow et al., 2018; Marisca et al., 2020). To do this *Tg(sox10:gal4); Tg(uas:gCaMP6s)* animals at 3 dpf were imaged. We first imaged baseline Ca^2+^ at 100ms for two minutes in sensory OLCs at 23 °C. These movies displayed full cell or sheath Ca^2+^ transients that could be detected in non-sensory OL but not sensory OLC (Fig. 7E) (n=9 animals total). These data demonstrate that in homeostatic conditions, sensory OLC do not display spontaneous Ca^2+^ transients like non-sensory OLCs. We may, at the least, expect Ca^2+^ transients to be detectable upon evoking the sensory circuit. To test this we exposed animals to 4°C then measured gCaMP6s intensity in both sensory and nonsensory OLCs. Nonsensory OLCs displayed an immediate spike in Ca^2+^ transient activity upon exposure to 4°C (Fig. 7F) (n=5 nonsensory OLCs). Alternatively, sensory OLCs did not display full cell/sheath Ca^2+^ transients after exposure to 4°C compared to 23°C (Fig. 7G) (n=4 sensory OLCs). The average number of peaks with a z score above 2 was significantly different from 2.0 ± 0.548 in nonsensory OLCs compared to 0.25 ± 0.25 in sensory OLCs (Fig. 7H) (n=9 animals). We then compared the average number of peaks in sensory OLCs with a z score above 2 in 23°C vs 4°C animals and there was no change and the z scores remained at an average of 0.25 ± 0.25 peaks above threshold, or a z score below 2 (Fig. 7I,G) (p>0.9999). These data demonstrate an additional district property of sensory OLCs from non-sensory OLCs.

### Sensory OLCs are important for somatosensory responses

Our results indicate that sensory OLCs are located at specific locations in the spinal cord at 5-10 and 13-18 DRGs (Fig. 1D). We, therefore, next considered the hypothesis that sensory OLCs may be required somehow in the sensory response. The DRG neurons fire upon immersion in 4°C, resulting in a hypothermic shivering behavior (Nichols and Smith, 2020; Kikel-Coury et al., 2021). To explore if sensory OLCs function in behavior, all detectable sensory OLCs on the right side of the animal were ablated at 3 dpf. Following ablation, animals were recovered for 24-hours. All animals that were tested displayed an intact sensory axon. Animals with ablated sensory OLCs were compared to animals with no ablations. As an additional control that the ablation laser itself did not elicit a change in behavior, the exact laser parameters used to ablate sensory OLCs were exposed to a region 10 μm from a sensory OLCs (n=24 animals total).

These animals were then immersed in 4°C for 10 sec and the percent of time shivering was calculated. Intact animals displayed a stereotypical shivering for 57.5% of time (intact vs. ablation control p=0.7803) (n=8 intact animals)(Fig. 8A,B). A similar phenotype was observed with ablation controls displaying shivering for 53.75% of time (intact vs. ablation control p=0.7803) (n=8 ablation control animals). This was in contrast to animals in which sensory OLCs were removed, which significantly reduced the shiver behavior to 2.5% of time spent shivering (intact vs. ablated sensory p<0.0001, ablation control vs. ablation sensory p<0.0001) (n=8 ablated sensory OLC animals) (Fig. 8A,B). To test if this eliminated other somatosensory responses, we performed an assay to determine an intact tactile response by touching the head of the animal with a pipette tip (Kikel-Coury et al., 2021) (Fig. 8C). Following this tactile assay, we could not detect differences in the percentage of intact, ablation control and sensory OLC ablated animals exhibiting a tactile response. 87.5% of intact control animals, 100% of ablation control animals, and 100% of ablated sensory OLC animals responded to the tactile assay (Fig. 8C) (n=24 animals total). In zebrafish at 4 dpf, Rohon Beard and trigeminal neurons detect tactile responses while the DRG do not (Ribera and Nuslein-Volhard, 1998). This indicates sensory OLCs could have a critical function in the DRG-specific behavioral circuit.

**Figure 8.**
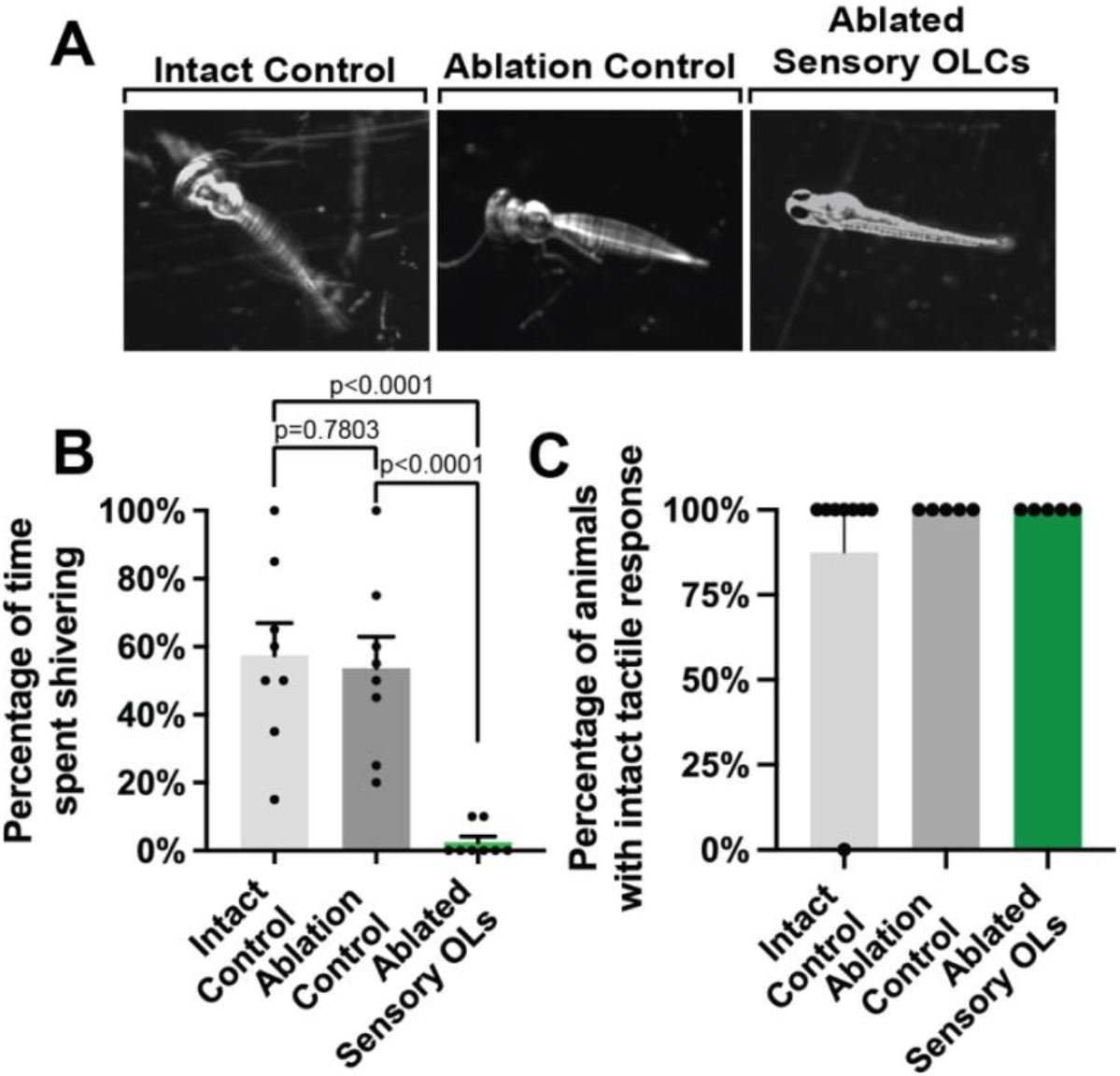
Sensory OLC ablation disrupts sensory behavior. (A) Images of three timepoints from a 20 second movies overlayed as a maximum projection in intact, ablation control, and ablated sensory OL animals showing the change in twitching behavior in sensory OL ablated animals upon exposure to cold water treatment. (B) Quantification of the percentage of time intact, ablated control, and sensory OL ablated animals spent shivering upon exposure to cold water treatment (p<0.0001, p=0.7803, p<0.0001). (C) Quantification of the percentage of intact, ablated, or sensory OLC ablated animals the exhibited a tactile response.

## DISCUSSION

Oligodendrocytes are essential for brain and spinal cord functionality (Nave and Werner, 2014; Allen and Lyons, 2018). In this study we demonstrate an OLC subpopulation that resides on the DRG axons in the spinal cord of zebrafish. These peripheral derived neurons provide a link between the CNS and PNS (Golding et al., 1997; Lumpkin and Caterina, 2007). Manipulations of DREZ genesis, where these DRG neurons cross from the periphery to the spinal cord, alters sensory-located OLCs. Artificial lesions that mimic a spinal cord axonal invasion supports the hypothesis that DREZ formation could be sufficient to impact these OLCs. Supporting a role of sensory OLCs in circuit function, ablation of sensory OLCs eliminates a DRG-specific somatosensory behavior (Nichols and Smith, 2020; Kikel-Coury et al., 2021). Together these data reveal a role for the DREZ in producing a unique OLCs population that functions in somatosensory neurons.

The traditional description of a mature oligodendrocyte is that it myelinates (Nave and Werner, 2014; Allen and Lyons, 2018). By the traditional definition, MBP-negative oligodendrocytes are simply immature. For this reason, we conservatively refer to the cells we uncovered as sensory OLCs instead of mature OLs. Although this is one interpretation, another is that MBP-negative oligodendrocytes are themselves mature oligodendrocytes with distinct functions. Pio del Rio Hortega proposed such a possibility, describing cells associated with blood vessels and neuronal somata as non-myelinating oligodendrocytes (Del Rio Hortega, 1911; Sierra et al., 2016). Single-cell sequencing of oligodendrocyte cells also demonstrate subpopulations of cells that express myelin markers at undetectable levels (Marques et al., 2016). Schwann cells, which myelinate the PNS, also present in non-myelinating mature forms (Jessen and Mirsky, 1997, 2005). Nonetheless, non-myelinating oligodendrocytes remain poorly understood. Our data supports the possibility that *mbp*^−^ OLC could be present in the spinal cord of zebrafish, albeit in low abundance and at very specific locations. In zebrafish, studies have shown heterogeneity of *mbp*^-^ OLCs with one population representing a subgroup that has less complexity like we visualize^5^. The potential functional role of sensory OLCs in a DRG-specific circuit within that circuit underscores their importance, regardless if they are a mature oligodendrocyte or an immature, long-lived progenitor.

The results above demonstrate the DREZ is important to produce a subpopulation of OLCs in the zebrafish of the spinal cord. Although our data supports the model that sensory and non-sensory OLCs are derived from a similar population of precursors cells in the spinal cord, our experiments do not distinguish between the possibilities that sensory OLCs are generated from a progenitor population that is unspecified until the DREZ forms or if a population is pre-determined to arrive to the DREZ once it is formed. It is clear, however, that sensory OLCs are derived from the ventral spinal cord region, likely the pMN domain. In mice, the presence of oligodendrocytes that are labeled with traditional myelin markers inhibits synaptic presence in ectopically located somatosensory axons in Semaphorin 6D mutants, suggesting that oligodendrocytes can inhibit synapses and thereby could be hypothesized to be required for somatosensory behaviors (Leslie et al., 2011).

Although our study reveals an interesting interaction between the DREZ and OLCs, it opens a gap in what molecular components at the DREZ drive such events. Discovering such molecules will help to further delineate myelinating oligodendrocytes from those that do not express typical myelin markers. Given that breach alone was sufficient to induce such a sensory OLC state, it seems possible the molecules released after injury could be involved. We also identified that sensory OLCs contact the DREZ, indicating that both secreted and short-range cues could be involved in the development of sensory OLCs. Studies that outline oligodendrocyte responses to injury will also be important guides for such discovery of the DREZ cue.

Our results also indicate an interesting change in sensory-induced behavior when sensory OLCs are reduced. This result was surprising given such a small number of ablated cells had such a profound impact on behavior. At the moment, our interpretation of that result is limited given the potential caveats of cell ablation in the intact spinal cord; although the control animals showed normal behavior. We also cannot explain why sensory OLCs appear at two areas of the spinal cord, especially given that spinal cord development should occur similarly on the anterior/posterior axis. However, the location of these cells in limited regions and the change in behavioral responses after their ablation could be linked. One possibility is that these cells function in synapse maturity of the sensory circuit and thus are located at regions where two neurons must be connected. It will be imperative to have a detailed and careful mapping of the sensory circuit in order to investigate such a possibility. Regardless, it will be important in the future to further investigate the presumed impact on somatosensory driven behaviors.

Our study establishes the role of PNS development in connectivity and function of the spinal cord, expanding the potential mechanisms that regulate OLC development. Collectively, the OLC heterogeneity in this study includes anatomical, structural, developmental and functional attributes.

## METHODS AND MATERIALS

### Ethics Statement

All animal studies were approved by the University of Notre Dame Institutional Animal Care and Use Committee.

### Contact for Reagent and Resource Sharing

Further information and requests for resources and reagents should be directed to the Lead Contact, Cody J. Smith. All of the data for this study is included in the figures.

### Experimental Model and Subject Details

All animal studies were approved by the University of Notre Dame IACUC as noted above. The following zebrafish strains were used in this study: AB, *Tg(sox10:mrfp* (Kucenas et al., 2008), *Tg(sox10:megfp* (Smith et al., 2014), *Tg(nkx2.2a:gfp* (Kifby et al., 2006), *Tg(sox10:eos* (McGraw et al., 2012), *Tg(sox10:gal4;uaslifeact-gfp*) (Helker et al., 2013; Hines et al., 2015), *Tg(gfap:nsfb-mcherry)* (Johnson et al., 2016; Smith et al., 2016), *Tg(olig2:dsred)* (Park et al., 2002b), *Tg(dbx:gfp)* (Briona and Dorsky, 2014), *Tg(nbt:dsred*(Peri and Nüsslein-Volhard, 2008), *Tg(ngn1:gfp)* (Prendergast et al., 2012), and *Tg(mbp:gfp)*(Czopka et al., 2013). All germline transgenic lines used in this study were stable unless otherwise noted. Pairwise matings were used to produce embryos and raised at 28°C in egg water in constant darkness. Animals were staged by hours or days post fertilization (hpf and dpf)(Kimmel et al., 1995). Embryos of either sex were used for all experiments.

### Method Details

#### *In vivo* imaging

Animals were anesthetized using 3-aminobenzoic acid ester (Tricaine), covered in 0.8% low-melting point agarose. Animals were then mounted laterally on their right side in glass-bottomed 35 mm petri dishes(Nichols et al., 2018). Images were acquired on a spinning disk confocal microscope custom built by 3i technology (Denver, CO) that contains: Zeiss Axio Observer Z1 Advanced Mariana Microscope, X-cite 120LED White Light LED System, filter cubes for GFP and mRFP, a motorized X,Y stage, piezo Z stage, 20X Air (0.50 NA), 63X (1.15NA), 40X (1.1NA) objectives, CSU-W1 T2 Spinning Disk Confocal Head (50 μm) with 1X camera adapter, and an iXon3 1Kx1K EMCCD camera, dichroic mirrors for 446, 515, 561, 405, 488, 561,640 excitation, laser stack with 405 nm, 445 nm, 488 nm, 561 nm and 637 nm with laser stack FiberSwitcher, photomanipulation from vector high speed point scanner ablations at diffraction limited capacity, Ablate Photoablation System (532 nm pulsed laser, pulse energy 60J @ 200 HZ). Images in time-lapse microscopy were collected every 5 min for 24 hours or from 24-72 hours depending on the experiment. Adobe Illustrator, ImageJ, and IMARIS were used to process images. Only brightness and contrast were adjusted and enhanced for images represented in this study. All fluorescence quantifications were normalized to the background value of each image.

#### Single-cell ablations

*Tg(sox10:mrfp)*, and *Tg(sox10:megfp)* animals were anesthetized using 0.02% 3-aminobenzoic acid ester (Tricaine) in egg water. Fish were then mounted in 0.8% low-melting point agarose solution, arranged laterally on a 10 mm glass-coverslip bottom petri dish, and placed on the microscope anterior to posterior. Confocal z-stack images of sensory and nonsensory OLCs in individual transgenic animals at 72 hpf were taken pre-ablation. Confirmed sensory and nonsensory cells were brought into a focused ablation window. Upon focusing the confirmed cell, we double-clicked on the center of the cell body using a 4 μm cursor tool to fire the laser(Nichols et al., 2018; Green et al., 2019). All laser parameters used are specific to our confocal microscope. Specific parameters include Laser Power (2), Raster Block Size (4), Double-Click Rectangle Size (8), and Double-Click Repetitions (4).

#### Single OL imaging over multiple days

Confocal z-stack images of *Tg(nkx2.2a:gfp);Tg(sox10:mrfp)* animals at 3 dpf were taken to locate and confirm the presence of sensory OLCs. After locating a sensory OLC, we counted DRGs anteriorly from the DRG most adjacent to the sensory OLC. We also counted the somites of the animal anteriorly from the somite number most adjacent to the location of the sensory OLC. At 4 dpf, we used the DRG number and somite number to locate the same sensory OLC within the same animal. Upon locating the sensory OLC we took a confocal z-stack image. We repeated the location and imaging process at 5 dpf and 6 dpf. We quantified the length and width of sheathes of individual sensory OLCs over a 4-day period using ImageJ.

#### Chemical treatments

The chemical treatments used for this study include SU6656 (Santa Cruz Biotechnology) and paclitaxel (Acros)(Nichols and Smith, 2019a). Stock solutions of both reagents were kept at − 20°C. SU6656 was kept at a concentration of 375 μm. Paclitaxel was kept at a concentration of 2.2 mM(Nichols and Smith, 2019a). All embryos were dechorionated at 36 hpf and incubated with 3 μM (SU6656) and 22 μM (paclitaxel) in egg water until imaging(Nichols and Smith, 2019a). All control animals were incubated with 1% DMSO in egg water.

#### Dorsal root ganglia (DRG) ablations

*Tg(sox10:mrfp);Tg(ngn1:gfp)* 48 hpf animals were anesthetized using 0.02% 3-aminobenzoic acid ester (Tricaine) in egg water. Fish were then mounted in 0.8% low-melting point agarose solution, arranged laterally on a 10 mm glass-coverslip bottom petri dish, and placed on the microscope anterior to posterior. Following the formation of pioneer axons from the DRG, we selected an 8 μm area between the DRG and the pioneer axon and double-clicked to fire the lesion laser(Nichols et al., 2018; Green et al., 2019). All laser parameters used are specific to our confocal microscope. Specific parameters include Laser Power (3), Raster Block Size (2), Double-Click Rectangle Size (8), and Double-Click Repetitions (4).

#### PA-Rac1

A *tol2–4xnr UAS-PA Rac1-mcherry* plasmid was diluted to 12 ng/μl in water along with 25 ng/μl of *transposase* mRNA. The injection solution was injected into *Tg(sox10:gal4); Tg(uas:lifeact-GFP)* embryos. Embryos were screened for *mcherry* expression via confocal microscopy at 48 hpf. Mature PA-Rac1 photoactivates upon exposure to 445 nm light. We imaged *mcherry^+^* DRGs for 24 h with exposure to 445 nm light. As a control we imaged *mcherry^+^* DRGs for 24 h without exposure to 445 nm light. *tol2–4xnr UAS-PA Rac1-mcherry* was a gift from Anna Huttenlocher (Addgene plasmid #41878)(Yoo et al., 2010).

#### Radial glia focal lesion

Focal lesions of the radial glia membrane to mimic a DREZ injury were created using the Ablate!TM© Photoablation System described above(Nichols and Smith, 2019a). Two laser induced lesions were created at 48 hpf. Each focal lesion ranged from 10-12 μm in size and was created 10 μm from the ventral radial glial membrane. This area best represented the location of the DREZ. Non-lesioned DREZ areas of adjacent nerves served as controls. Following laser induced focal lesioning, time-lapse images of the nerves were taken as described above for 24 hours.

#### Immunohistochemistry

The primary antibody used to determine mature myelination was anti-mbp (1:500, rabbit, Appel Lab)(Kucenas et al., 2009). The secondary antibody used was Alexa Fluor 647 goat antirabbit (1:600; Invitrogen). 15 dpf animals were fixed in 4% PFA in PBST (PBS, 0.1% Triton X-100) at 25°C for 3 hours. Fixed animals were washed with PBST, DWT (dH_2_0, 0.1% Triton X-100), and acetone for 5 minutes each. Next, the animals were incubated in −30°C acetone for 10 minutes and then washed three times with PBST for 5 minutes each. Animals were then incubated with 5% goat serum in PBST for 1 hour at 25°C, then, the primary antibody solution was added for 1 hour at 25°C. After 1 hour, then animals were transferred to 4°C overnight. Fixed animals were washed three times with PBST for 30 minutes each and once more for 1 hour. Animals were then incubated with secondary antibody solution for 1 hour at 25°C, then transferred to 4°C overnight. Fixed animals were washed three times with PBST for 1 hour and immediately imaged. Animals were mounted, and confocal images were taken using the protocol above for in vivo imaging.

#### Calcium Imaging

*Tg(sox10:gal4); Tg(uas:gCaMP6s)* and *Tg(neurod:gal4);Tg(uas:gCaMP6s)* 3 dpf animals were mounted in 0.6% low-melting point agarose solution, arranged laterally on a 10 mm glass-coverslip bottom petri dish, and placed on the microscope anterior to posterior. Animals were not anesthetized. For neuronal imaging, spinal cord z-stacks were collected that spanned 10 μm (1 μm steps) and imaged for 3 min with 2 second intervals. For OLC imaging, confocal z-stack images of the spinal cord in individual transgenic animals at 3 dpf were taken to confirm the presence of sensory and nonsensory OLCs. Upon confirmation, sensory OLCs were imaged every 2 seconds for 2 minutes to capture Ca^2+^ transients in room temperature (23 degree) egg water. After imaging baseline Ca^2+^ activity, room temperature egg water was removed and 3 mL of 4 degree cold egg water was added. Animals were then imaged immediately following the addition of 4 degree egg water and after refocusing. Nonsensory OLCs were also imaged every 2 seconds for 2 minutes as a control. The exposure was set to 100 ms.

#### Animal behavior after sensory OLC ablations

*Tg(sox10:mrfp)* animals at 72 hpf were anesthetized using 0.02% 3-aminobenzoic acid ester (Tricaine) in egg water. Fish were then mounted in 0.8% low-melting point agarose solution, arranged laterally on a 10 mm glass-coverslip bottom petri dish, and placed on the microscope anterior to posterior. Confocal z-stack images of *Tg(sox10:mrfp)* animals at 72 hpf were taken to locate and confirm the presence of sensory OLCs. We ablated sensory OLCs using the Ablate!TM© Photoablation Laser following the single-cell ablation methods described above. Following successful ablation, we unmounted individual animals from the agarose solution. We then placed individual animals into a small petri dish containing 4°C egg water and recorded their twitching response to cold water treatment for a 20 second period. As a control for the ablation, we ablated a small 4 μm region above the sensory axon where a typical sensory OL would reside. As another control, we imaged separate set of intact control animals that had unablated sensory OLC. The ablated, ablated control, and intact control animals were each exposed to 4°C egg water and their twitching response to cold water treatment was recorded over a 20 second period.

### Quantitative Analysis

#### Quantification and statistical analysis

To generate composite z-images for the cell, 3i Slidebook software (Denver, CO) was used. Individual z-images were sequentially observed to confirm composite accuracy. All graphically presented data represent the mean of the analyzed data unless otherwise noted. Cell tracking was performed using the MTrackJ plugin for ImageJ (https://imagescience.org/meijering/software/mtrackj/, Bethesda, MD). GraphPad Prism software (San Diego, CA) was used to determine statistical analysis. Full detail of the statistical values can be found in <u>S1 Table</u>.

#### Quantification of Direction Changes

To track the direction changes of OLCs, we imaged *Tg(sox10:mrfp)* and *Tg(sox10:gal4:uas:lifeact-gfp)* animals from 48-72 hpf. We then tracked single *sox10*^+^ OLCs in each animals throughout the 24 time-lapse movie using the MTrackJ plugin of ImageJ. We quantified the total distance an individual OLC traveled as well as the amount of direction changes an individual OLC experienced throughout the 24-hour imaging window.

#### Quantification of migration

To track the individual paths of sensory and nonsensory OLCs, the MTrackJ plugin on ImageJ was used. The center of each cell body was traced over time, and quantitative data were collected by ImageJ. Resulting x and y coordinates of each cell were overlaid to create migration plots. All distance and time points were calculated by the MTrackJ software and further quantified using Microsoft Excel (Redmond, WA).

#### Quantification of sheath length and width

To determine the length and width of sensory and nonsensory OL sheaths, we used the freehand tracing tool in ImageJ. We traced the ensheathed area longitudinally and laterally and used the Analyze feature within ImageJ to obtain quantitative results.

#### Quantification of velocity

To obtain the velocity of an individual OLC traveling, we tracked individual paths of sensory and nonsensory OLCs throughout the timelapse imaging window using the MTrackJ plugin on ImageJ. We tracked the center of each cell body over time and used the quantitative distance and time collected by ImageJ to calculate velocity. We calculated the velocity using Microsoft Excel (Redmond, WA).

#### Quantification of calcium imaging

We imaged *Tg(sox10:gal4);Tg(uas:gCaMP6s)* 3 dpf animals every 2 seconds for 2 minutes to visualize Ca^2+^ transient waves in sensory and nonsesory OLCs exposed to 4°C and 23°C degree egg water. Later, we enlarged the movies to visualize one sensory or nonsensory OLC at a time. We used the line tool in ImageJ to trace individual projections of OLCs and then measured the integrated density along the projection using the integrated density tool within ImageJ. In order to assess changes in integrated density, we then used Microsoft Excel to calculate the Z score of the integrated density for each projection. Any Z score > or = 2 were noted as positive for Ca^2+^ transient activity. Any Z score < 2 were not recognized as positive for Ca^2+^ transient activity. We first tested this by imaging *Tg(neurod:gal4); Tg(uas:gCaMP6s)* and measured calcium transients in DRG neurons after exposure to 4°C or 23°C water. We imaged *Tg(neurod:gal4): Tg(uas:gCaMP6s)* 3dpf animals every 2 seconds for 3 minutes to assess changes in gCaMP6s fluorescence of DRG neurons in response to 23°C and 4°C egg water. Movies were analyzed in ImageJ by tracing individual DRG neuron somas and measuring integrated density for each timepoint. In this analysis, the Z score of DRG neuron integral densities were measured and used to assess changes in integral density. If gCaMP6s integral density z score was 2 or greater, this was marked as an active firing event.

#### Quantification of animal behavior

We recorded the behavior of sensory ablated, ablation control, and intact animals for 20 seconds immediately following exposure to 4°C egg water. Any movies longer than 20 seconds were normalized to 20 seconds. We recorded the amount of time each animal spent shivering in cold water treatment. We divided the total time an animal spent shivering over the total time recorded. All calculations were performed using Microsoft Excel (Redmond, WA).

## ACKNOWLEDGEMENTS

We thank David Lyons, Bruce Appel and Ethan Scott for transgenic zebrafish lines, Bruce Appel, Jacob Hines, and members of the Smith and Wingert labs for their helpful comments. We would also like to thank Sara Cole and the Notre Dame Imaging Core for their help using IMARIS software. This work was funded by the Alfred P. Sloan Foundation (CJS) and the NIH (DP2NS117177). Further support was provided by the University of Notre Dame, the Elizabeth and Michael Gallagher Family, Center for Zebrafish Research at the University of Notre and Center of Stem Cells and Regenerative Medicine at the University of Notre Dame.

## CONTRIBUTIONS

LAG, RG, JB and ELN conducted all experiments. LAG, RG, JB and CJS analyzed all experiments. LAG and CJS wrote the manuscript with input from RG and JB. CJS and LAG conceived the project.

## SUPPLEMENTARY INFORMATION

**Table S1.**
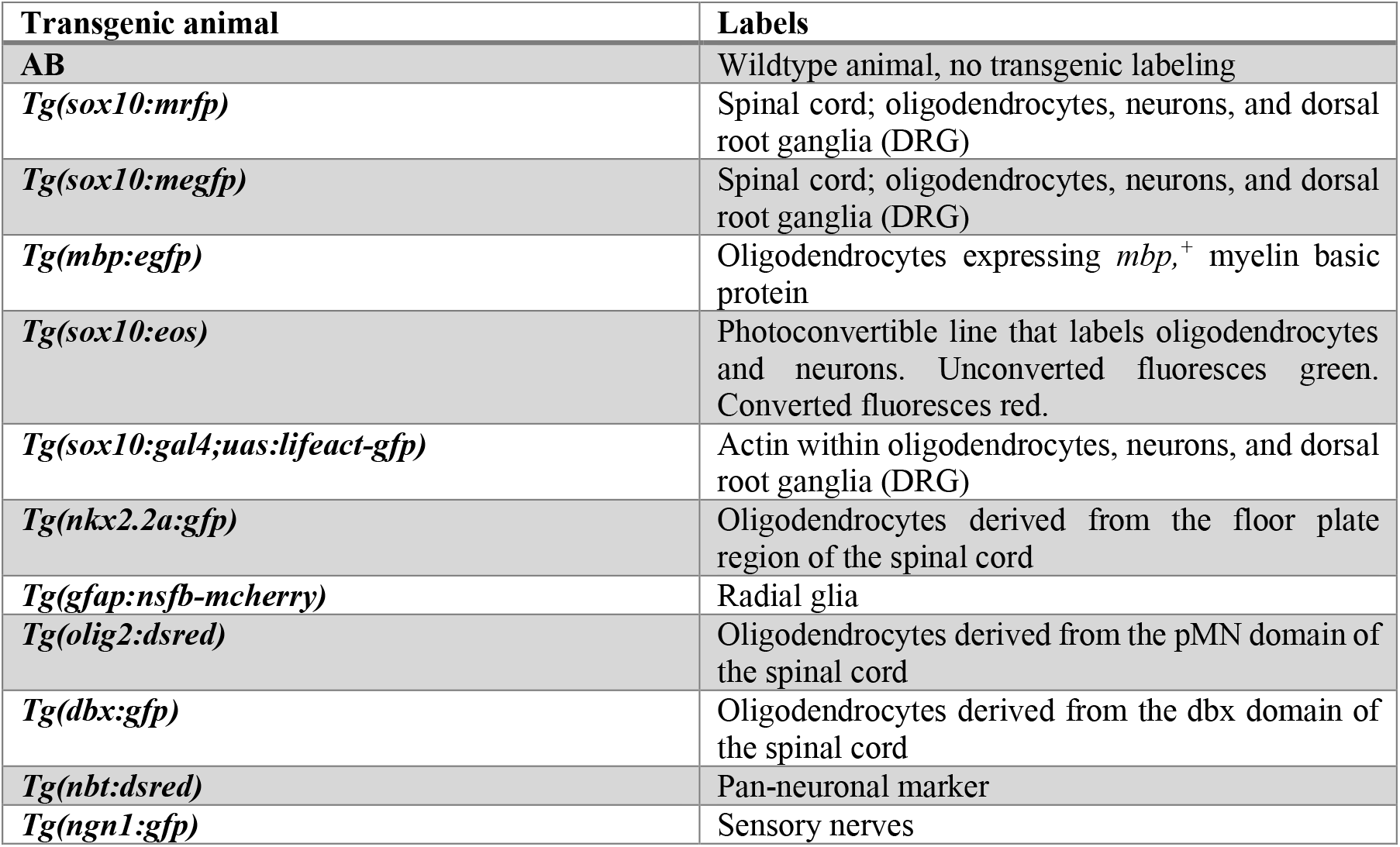
List of the transgenic zebrafish strains used in the study.

**Table S2. Statistics Sheet.** All raw values of the individual cell and zebrafish n-values scored for each figure. Table also includes specific values for statistical tests and exact p-values used to determine statistical significance for each figure panel.

**Figure S1.**
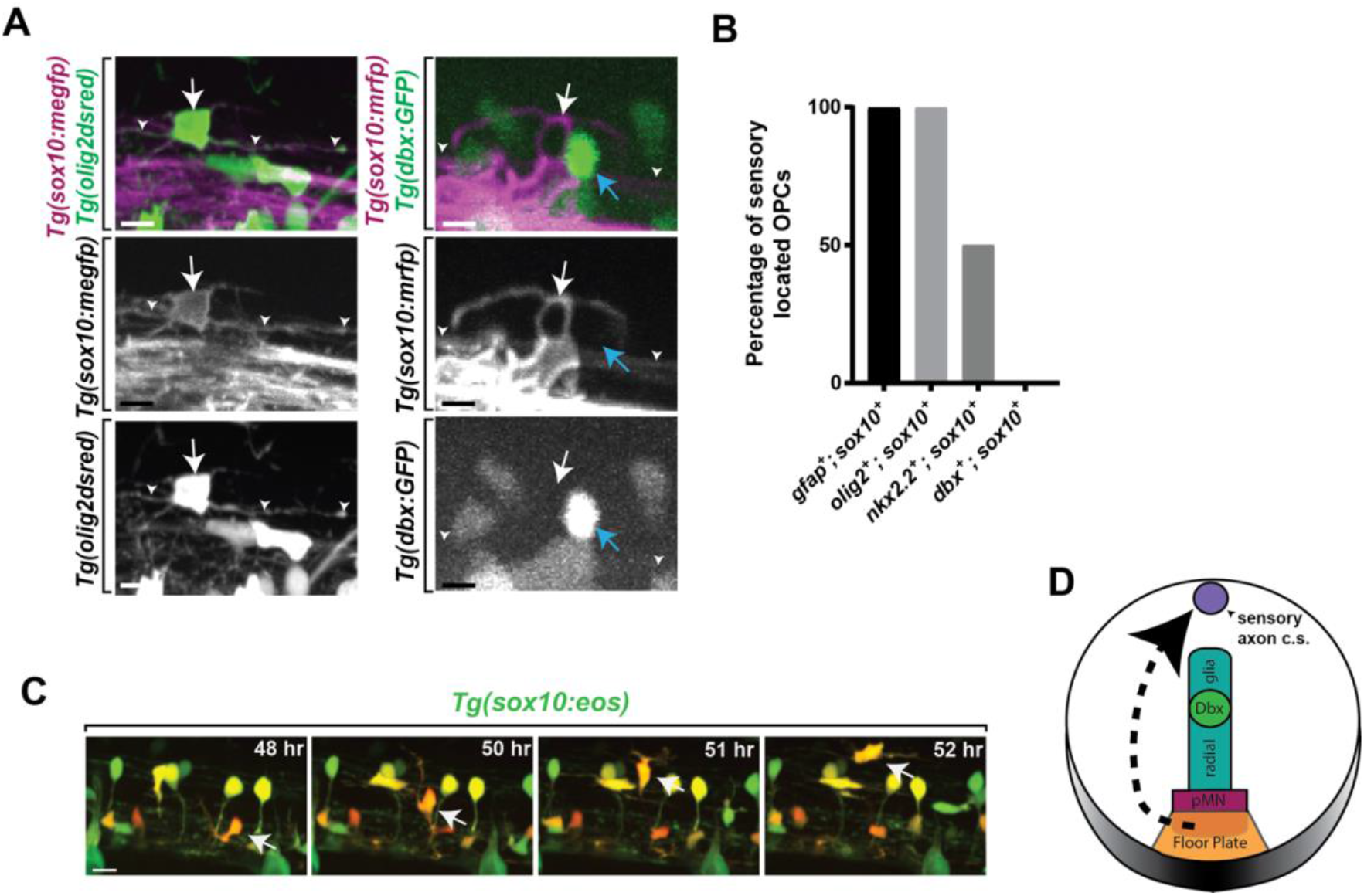
Sensory oligodendrocytes are not generated from a distinct progenitor pool. (A) Confocal z-stack still images taken at 72 hpf in *Tg(sox10:megfp):Tg(olig2:dsred)*, and *Tg(sox10:mrfp);Tg(dbx:gfp)* animals showing the presence or absence of progenitor marker expression in sensory OLs. White arrows represent sensory OLs. Blue arrow represents nonsensory OL. White arrowheads represent the sensory axon. (B) Quantification of the percentage of sensory located oligodendrocytes that are *olig2^+^;sox10^+^*, *nkx2.2a^+^;sox10^+^, or dbx^+^;sox10.^+^* (C) Images from a 24 hour time-lapse starting at 48 hpf in *Tg(sox10:eos)* animals showing the migration of a photoconverted OPC from the ventral spinal cord to the dorsal spinal cord region. White arrow represents a photoconverted OPC that migrates to the sensory. (D) Schematic diagram depicting the migration of sensory oligodendrocytes from the floor plate region *(nkx2.2a)* to the sensory axon. Scale bar equals 10μm (A,C).

**Figure S2.**
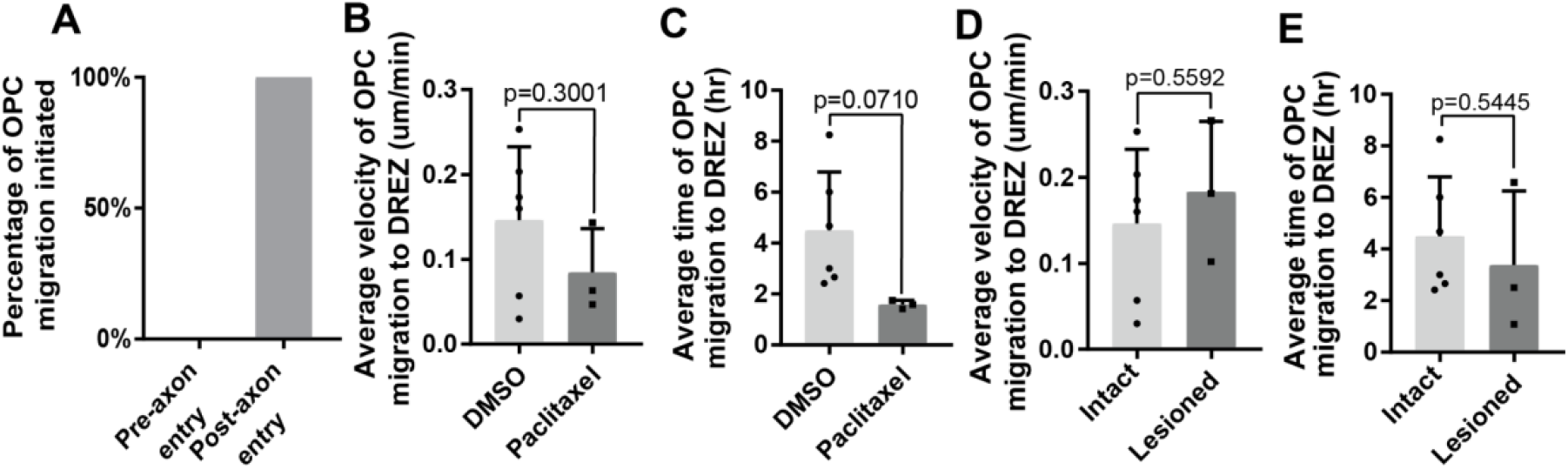
Sensory OLCs arrive following axon entry and DREZ contact. (A) Quantification of the percentage of OPC migration initiated showing OPC migration occurs post-axon entry. (B) Quantification of the average velocity of OPC initiation to the DREZ in DMSO and Paclitaxel treated animals (p=0.3001). (C) Quantification of the average time of OPC migration initiation to the DREZ in DMSO and Paclitaxel treated animals (p=0.0710). (D) Quantification of the average velocity of OPC initiation to the DREZ in intact and lesioned treated animals (p=0.5592). (E) Quantification of the average time of OPC migration initiation to the DREZ in intact and lesioned treated animals (p=0.5445).

